# Towards a global barcode reference library for subterranean fauna

**DOI:** 10.1101/2024.09.20.613981

**Authors:** Michelle T. Guzik, Jake Thornhill, Mieke van der Heyde, Danielle N. Stringer, Nicole E. White, Mattia Saccò, Perry G. Beasley-Hall, Paul Nevill, Rachael A. King, Steven J. B. Cooper, Andrew D. Austin

## Abstract

Implementation of environmental DNA (eDNA) metabarcoding for biodiversity discovery and assessment offers a unique opportunity to gain new insights into subterranean communities around the world. However, for effective and meaningful identification of species from anonymous eDNA barcodes, a library of known reference sequences with associated correct taxonomic metadata -also called a barcode reference library (BRL) - is required. Here we propose an open, publicly accessible information resource for eDNA biomonitoring of subterranean fauna following findable, accessible, interoperable and reusable (FAIR) principles that can be expanded globally. While similar proposals have been made by other authors for individual taxon groups, here we have analysed a curated BRL of subterranean fauna compiled from existing GenBank and BOLD DNA sequence databases for a minimum of four genes (*COI, 18S*, *12S*, and *16S* rRNA). We demonstrate the value of such an initiative to eDNA metabarcoding where custom libraries are used to characterise entire ecosystems under examination. To examine the effectiveness of a custom BRL, we generated metabarcoding data for an exemplar system at Bungaroo Creek in the Pilbara (Australia), a globally significant location of stygofaunal diversity. We compared results of BLAST queries of eDNA Operational Taxonomic Units (OTU’s) to our BRL and the GenBank nucleotide database, and observed that for all barcoding regions, the custom BRL identified subterranean ZOTUs that could not be identified using GenBank and worked better in tandem. We use the BRL presented here to propose a four-stage plan for developing data infrastructure for subterranean fauna, especially with respect to eDNA metabarcoding data. To our knowledge, this represents the first published instance of a subterranean BRL being tested against real eDNA metabarcoding data. These findings provide a step forward towards robust DNA-based bioassessments for subterranean biodiversity and further emphasise the need for the eDNA community to work together in facilitating a global BRL.

## Introduction

Water health underground is naturally maintained through purification and bioremediation services performed by organisms associated with groundwater aquifers (Boulton *et al*. 2008; Griebler and Avramov 2015). The total composition of these biological communities is largely unknown, but the macro-eukaryotic fauna that occupy scavenging, filtering and predatory roles are generally a better studied component of groundwater-dependent ecosystems (Koch *et al*. 2024). Collectively described as subterranean fauna or stygofauna, these animals include fish and invertebrates (from multiple phyla and orders) that characteristically lack eyes and body pigment, and frequently have exaggerated body shapes and sensory appendages. Survivors of millions of years of global climatic and biogeographic change (Page *et al*. 2008), these ecosystems represent ancient evolutionary “time capsules” that are frequently restricted to individual aquifers (Guzik *et al*. 2008), with most species therein being undescribed (Guzik *et al*. 2011a). Given the current and future extent of human impacts on groundwater environments and their biodiversity worldwide (Sala 2000; Mammola *et al*. 2024), the need for accelerated identification tools and bioassessment of these understudied, hyperdiverse (Meierotto *et al*. 2019), and threatened ecological communities (Butchart *et al*. 2010) is especially urgent (Mammola *et al*. 2019; Mammola *et al*. 2024).

Traditional biomonitoring methods typically require targeted sampling, species identification using morphology and confirmatory single-gene DNA sequencing (Beng and Corlett 2020). High-throughput sequencing has created a revolutionary opportunity for a cheaper (Wang et al. 2018), less invasive (Beja-Pereira *et al*. 2009), and potentially more accurate (Lejzerowicz *et al*. 2015) addendum to biological assessment and documentation of biodiversity through the metabarcoding analysis of environmental DNA (eDNA) (Ficetola *et al*. 2008; Bohmann *et al*. 2014; Cristescu 2014; Bista *et al*. 2017; Takahashi *et al*. 2023). However, to date only a few studies have focused on using eDNA to assess and monitor species assemblages in groundwater (Goricki *et al*. 2016; Korbel *et al*. 2017; Niemiller *et al*. 2018; Couton *et al*. 2023a; Couton *et al*. 2023b; van der Heyde *et al*. 2023a; van der Heyde *et al*. 2023b). A key reason for low uptake of this method is the difficulty in identifying anonymous eDNA operational taxonomic units (OTUs) solely using the online GenBank repository as a reference database which often does not provide a high level of taxonomic resolution for such data, especially for poorly known taxonomic groups which typify subterranean communities (see Box 1 for additional information).

### Box 1

#### OTUs, databases and predicaments for subterranean fauna

As a consensus of raw eDNA metabarcoding reads, OTUs are estimated bioinformatically using a predetermined threshold (Rideout *et al*. 2014) and represent the DNA signature of organisms present in an original environmental sample. Likewise, each OTU acts as a ‘species’ proxy (Marques *et al*. 2020) that can be used to infer estimates of biodiversity. By nature, OTUs are anonymous, lacking associated metadata other than locality information for the eDNA sample. The absence of associated taxonomic information in eDNA reads is problematic for biomonitoring because diagnosis of specimens to the lowest taxonomic rank (i.e. species level) is required if estimates of biodiversity are to be accurate and conservation status listings are to be enacted (Belle *et al*. 2019; Beng and Corlett 2020). In order to attribute taxonomic information to OTUs, it is essential to query each sequence against accurate and detailed sequence databases that contain associated taxonomic metadata (Boykin *et al*. 2012; Coissac *et al*. 2012; Kress *et al*. 2015; Rees *et al*. 2015; Bissett *et al*. 2016; Creer *et al*. 2016). Currently, the most readily queried genetic sequence databases for taxonomic identification of OTUs are the National Center for Biotechnology Information’s (NCBI) GenBank repository and the International Barcode of Life Consortium’s (iBOL) Barcode of Life Database (BOLD).

Both GenBank and BOLD are a valuable resource for identification of common species but have numerous limitations for querying OTUs where subterranean fauna is concerned. In particular, there is a limited availability of sequence data (i.e. many unknown species, sometimes representative of higher-level taxa, i.e. genera, tribes, etc) and an inaccuracy in taxonomic identifications (Tixier *et al*. 2012), despite taxonomy-free approaches being developed (Wilkinson *et al*. 2024; Mächler *et al*. 2021b). Due to the voluntary submission of data to public databases, only 15–25% of described animal species are represented on GenBank (Kvist 2013; Dormontt *et al*. 2018; DeWaard *et al*. 2019) and less so on BOLD, which has an historical bias towards characterisation of the *cytochrome c oxidase subunit I* (*COI*) gene (Hebert *et al*. 2003; Hebert and Gregory 2005). Further, a significant proportion of the barcode records in these reference databases are only identified to family level (Kwong *et al*. 2012; Curry *et al*. 2018), in part because some 90% of multicellular species are undescribed (DeWaard *et al*. 2019). The problem is further exacerbated by the fact that numerous species in databases are likely misidentified (e.g. Bridge *et al*. 2003; Vilgalys 2003; Meiklejohn *et al*. 2019). Whilst BOLD requires voucher specimens, this is not the case for GenBank, so potential misidentifications can never be checked. Whilst publicly available libraries with comprehensive species coverage are an internationally recognised goal that will facilitate genetic diversity and promote new species identification (Ratnasingham and Hebert 2007; Zahiri *et al*. 2017), all the existing public DNA sequence databases are still extremely limited in depth of biodiversity and taxonomic metadata. Customised barcode reference libraries (BRL) can improve resolution when assigning OTUs especially when unknown biodiversity in communities is high and groundwater ecosystems are no exception. For identification of OTUs to taxonomic levels beyond order (van der Heyde *et al*. 2023b) and for eDNA metabarcoding to be successfully implemented in biomonitoring of subterranean fauna, a global BRL is essential.

Given the breadth and depth of DNA sequence data and its associated metadata currently being collected in biological studies, reusability and accessibility are broadly recognised as urgent priorities (Hassenrück *et al*. 2021). The scientific community largely agrees that the guiding principles of data management and infrastructure are that it be findable, accessible, interoperable and reusable (FAIR) (Wilkinson *et al*. 2016). However, in the non-academic sector, DNA sequences are known to be disparately databased and frequently held privately (Berry *et al*. 2021). This predicament also exists for subterranean fauna where substantial DNA sequence data are collected daily for species identification for legislative requirements (Gibson *et al*. 2019). However, commercial interest in maintaining siloed and private databases of subterranean fauna sequences exists prohibiting effective data sharing and FAIR and public BRLs.

### Global BRL database for subterranean fauna

There is a clear need to establish a ‘living’ dataset for eDNA biomonitoring of subterranean fauna that comprises a phylogenetic backbone (common ancestors) to which tips (groups of descendent species and/or individuals) can be added (Saclier *et al*. 2024). This generates an accurate representation of phylogenetic relatedness and recent genetic diversity which can be used as a molecular proxy for biodiversity (Floyd *et al*. 2002; Blaxter and Floyd 2003) and can be especially important for identification of cryptic and unknown species. The overall aim of the current study was to establish a subterranean BRL that follows the FAIR principles, meaning it is publicly available (findable, accessible) and a place where sequence representation can grow as samples are added, permits novel ways to analyse and use the data are developed, and new users can contribute to the resource (interoperable and reusable). Similar proposals have been made previously for specific taxonomic groups (pseudoscorpions (Hlebec *et al*. 2023); isopods and myriapods (Rendoš *et al*. 2023); asellid isopods (Saclier *et al*. 2024)). However, we have significantly expanded on these studies by curating sequences across the subterranean fauna tree of life, thus implementing the recommendation of Saccò *et al*. (2022b) and Guzik *et al*. (2024) and taking the first step towards developing an all-encompassing subterranean fauna BRL.

To demonstrate the possibilities for such a BRL database, especially for OTU queries derived from eDNA metabarcoding, we used the Pilbara region of Western Australia as an exemplar location, known for its extremely rich diversity of subterranean fauna and recognised internationally as a subterranean biodiversity hotspot (Figure 1). We took advantage of BOLD to facilitate curation of our BRL. While acknowledging that historically, BOLD focussed on the cytochrome c oxidase subunit I (*COI*) gene (Hebert *et al*. 2003), the platform currently accommodates data from multiple loci and hosts large taxonomic/biodiversity-based barcoding projects (Thomsen and Willerslev 2015); (reviewed by Zhou *et al*. 2009; Webb *et al*. 2012; Kvist 2013; Deagle *et al*. 2014; Wirta *et al*. 2016; Kuzmina *et al*. 2017; Aizpurua *et al*. 2018). BOLD has the advantage of an accessible browser-based interface which includes taxonomic, geographic, and photographic metadata for each sequence (Ratnasingham and Hebert 2007). Additionally, BOLD has a unified nomenclature for genetic lineages. Given nomenclatural and identification inconsistencies between projects (Meier *et al*. 2022) and the substantial taxonomic impediment affecting subterranean fauna (Mammola *et al*. 2021), BINs can be an effective option in unifying the nomenclature of ‘dark’ taxa in the absence of formal species descriptions.

**Figure 1:**
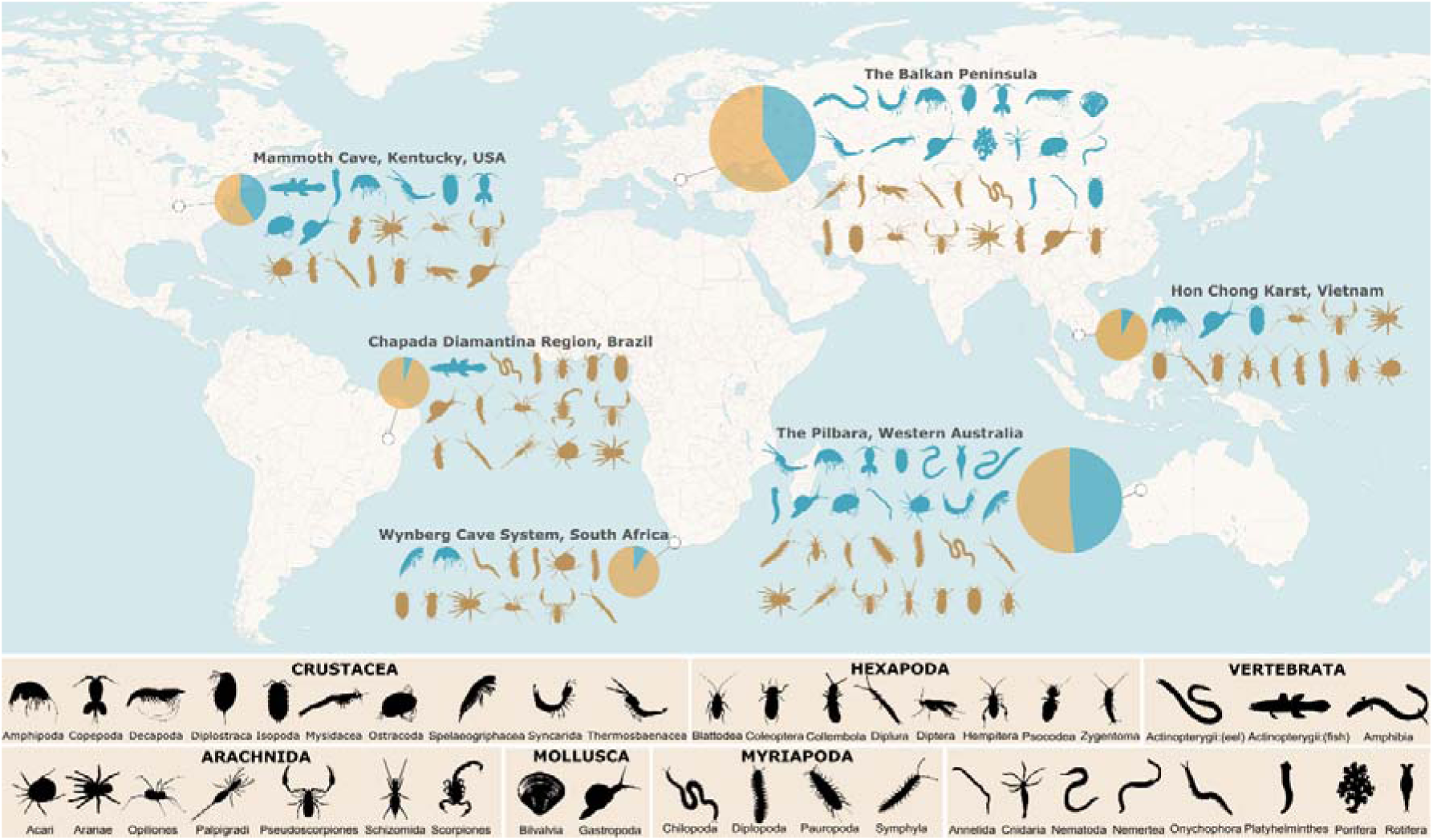
Global map showing single exemplar subterranean biodiversity hotspot locations per continent and respective major taxon group diversity of troglofauna (beige silhouettes) and stygofauna (blue silhouettes) (https://www.phylopic.org/) and the proportion of representative taxa per location (South America: Chapada Diamantina Region, Brazil (Gallão *et al*. 2023); North America: Mammoth Cave, Kentucky, U.S.A. (Niemiller *et al*. 2021); Europe: Balkan Peninsula (Sket *et al*. 2004); Asia: Hon Chong karst, Vietnam (Deharveng *et al*. 2023); Australia: Pilbara region, Western Australia (Halse 2018) and Africa: Wynberg cave system South Africa (Ferreira *et al*. 2020) with pie chart sizes: small = <100 subfauna species, large = >1,000. One hotspot per continent was chosen for conciseness to illustrate how the Pilbara compares to other well-known hotspots. See Supplementary Material 1 for species numbers. Map created in QGIS (2024) with base map layer: CartoDB, under CC BY 3.0. Data by OpenStreetMap, under ODbL.

Here we aimed to; (1) establish a preliminary phylogenetic backbone from existing published data from GenBank and BOLD data for six barcode genes, (2) test the applicability and validity of such a database by comparing BLAST queries of eDNA ZOTUs against two databases, GenBank and the custom BRL; and (3) discuss the areas that need to be addressed to facilitate the widespread implementation of the platform by the eDNA community (i.e. FAIR). Using this approach to querying subterranean eDNA data, we provide a baseline for researchers wishing to enhance the broader use of their data holdings, and provide a critical roadmap for future development of a global custom BRL for subterranean fauna.

## Methods

### Pilbara, Western Australia: a case study

The Pilbara region of Western Australia (WA) supports exceptionally diverse subterranean faunal communities first described in 1998 (Poore and Humphreys 1998) (Figure 1 and Supplementary Material 1). Well-known for its economically valuable natural resources (e.g., petroleum, off-shore natural gas, and mainland iron ore deposits), the Pilbara subterranean assemblages include blind vertebrates, such as the cave eel (Moore *et al*. 2018)) and gudgeon (Larson *et al*. 2013), and macroinvertebrates from multiple phyla and orders (e.g., (Eberhard *et al*. 2005; Guzik *et al*. 2011a; Guzik *et al*. 2011b; Karanovic and Cooper 2011; Karanovic and Cooper 2012; Halse *et al*. 2014; Karanovic *et al*. 2015; Perina *et al*. 2018; Mokany *et al*. 2019a; Mokany *et al*. 2019b; Perina *et al*. 2019; Matthews *et al*. 2020; King *et al*. 2022; Stringer *et al*. 2022)) that are specifically protected under the Australian Commonwealth Environment Protection and Biodiversity Conservation (EPBC) Act (1999) (Department of Climate Change, Energy, the Environment and Water 1999) and state-based Western Australian Biodiversity Conservation Act (2016) and the Environmental Protection Act (1986) (Environmental Protection Authority 2021). Five major river catchments are recognised in the Pilbara: Ashburton, Onslow Coast, Lyndon-Minilya, Port Hedland, and De Grey. They all contain areas of geology likely to comprise suitable habitat for subterranean fauna, namely unconsolidated sedimentary aquifers, including those in recent valley-fill alluvium and colluvium, coastal deposits, aquifers in chemically deposited calcretes and pisolites within Tertiary drainage channels, and fractured-rock aquifers in dolomites (Johnson and Wright 2001; Danielopol *et al*. 2003; Halse *et al*. 2014). The need for biomonitoring and assessment in these locations is as high in the Pilbara as anywhere else in the world, if not higher due to pressures from industrial mining.

### Pilbara subterranean barcode reference library curation

In order to build a preliminary BRL for subterranean taxa in the Pilbara, we curated sequences from GenBank and BOLD. Sequences were identified as suitable for inclusion based on a review of the literature (Supplementary Material 2) to identify studies that generated sequence data as PopSets (a collection of related DNA sequences derived from population, phylogenetic, mutation and ecosystem studies that have been submitted to GenBank) based on title, taxon groups known to contain subterranean fauna (GenBank), and consultants (specifically Biologic Environmental Survey Pty Ltd (Supplementary Material 2)) known to publish subterranean faunal data. Search terms included: Pilbara; subterranean fauna; Biologic; and authors known to publish on subterranean fauna taxon groups and key groups that contain subterranean fauna (Jarman and Elliot 2000; Leys *et al*. 2003; Cooper *et al*. 2007; Finston *et al*. 2007; Page *et al*. 2007; Cooper *et al*. 2008; Guzik *et al*. 2008; Leys and Watts 2008; Page *et al*. 2008; Guzik *et al*. 2009; Bradford *et al*. 2010; Chakrabarty 2010; Abrams *et al*. 2012; Asmyhr and Cooper 2012; Cook *et al*. 2012; Karanovic and Cooper 2012; Murphy *et al*. 2012; Abrams *et al*. 2013; Larson *et al*. 2013; Murphy *et al*. 2013; Asmyhr *et al*. 2014; Javidkar *et al*. 2015; Karanovic *et al*. 2015; Camacho *et al*. 2018; Moore *et al*. 2018; Matthews *et al*. 2020). We included trap door spiders (Araneae) in the search terms. Whilst these are not strictly troglofauna they are extremely important as short-range endemic fauna living in subterranean habitats in the Pilbara (Harvey *et al*. 2012).

The complete Pilbara subterranean fauna database was imported into DS-SUBFAUNA via BOLD (https://boldsystems.org/index.php/MAS_Management_DataConsole?codes=DS-SUBFAUNA). For a visual representation of the sequence data, a circular cladogram was reconstructed using the *Taxon ID Tree* function using the neighbour-joining algorithm (Saitou and Nei 1987) under a Kimura 2-parameter model (Kimura 1980). The BOLD Aligner (amino acid based hidden Markov model [HMM]) was used to align the sequences with default settings (Ratnasingham and Hebert 2007). As part of the BOLD system, BINs were assigned to all sequences. The BIN system is an interim taxonomic system, which clusters sequences using well established algorithms to produce operational taxonomic units that closely correspond to species (Ratnasingham and Hebert 2013).

### Groundwater eDNA sample collection

We also undertook field work at select locations to supplement published data. Twenty-three bores were sampled between June 2017 and October 2021 within Bungaroo Creek, Jimmawurrada Creek, and the Robe River alluvial aquifers of the Robe River Catchment (Bungaroo Study Area) for the presence of subterranean fauna eDNA (Figure 2) (Supplementary Material 3). A bailer was lowered down collared or cased boreholes to obtain five 1 L groundwater samples. All groundwater filtering equipment was decontaminated between sampling locations (i.e., individual boreholes) using 50% bleach solution and soaked for a minimum of 10 min. Once obtained, groundwater samples were transferred into sterile containers and refrigerated prior to filtering at a temporary field laboratory. Within 24 h of collection, groundwater was filtered using two Sentino peristaltic pumps (Pall Life Sciences, New York, USA). Each 1 L groundwater sample was passed through sterile cellulose filter membranes of either 0.22 µm or 0.45 µm pore size depending on the turbidity of the water. Individual filter membranes were folded in half, stored in separate sample bags, and frozen at −20° C for up to one month. Environmental control samples were obtained at open surface water points of the Robe River, filtered, and stored in the same manner as above.

**Figure 2:**
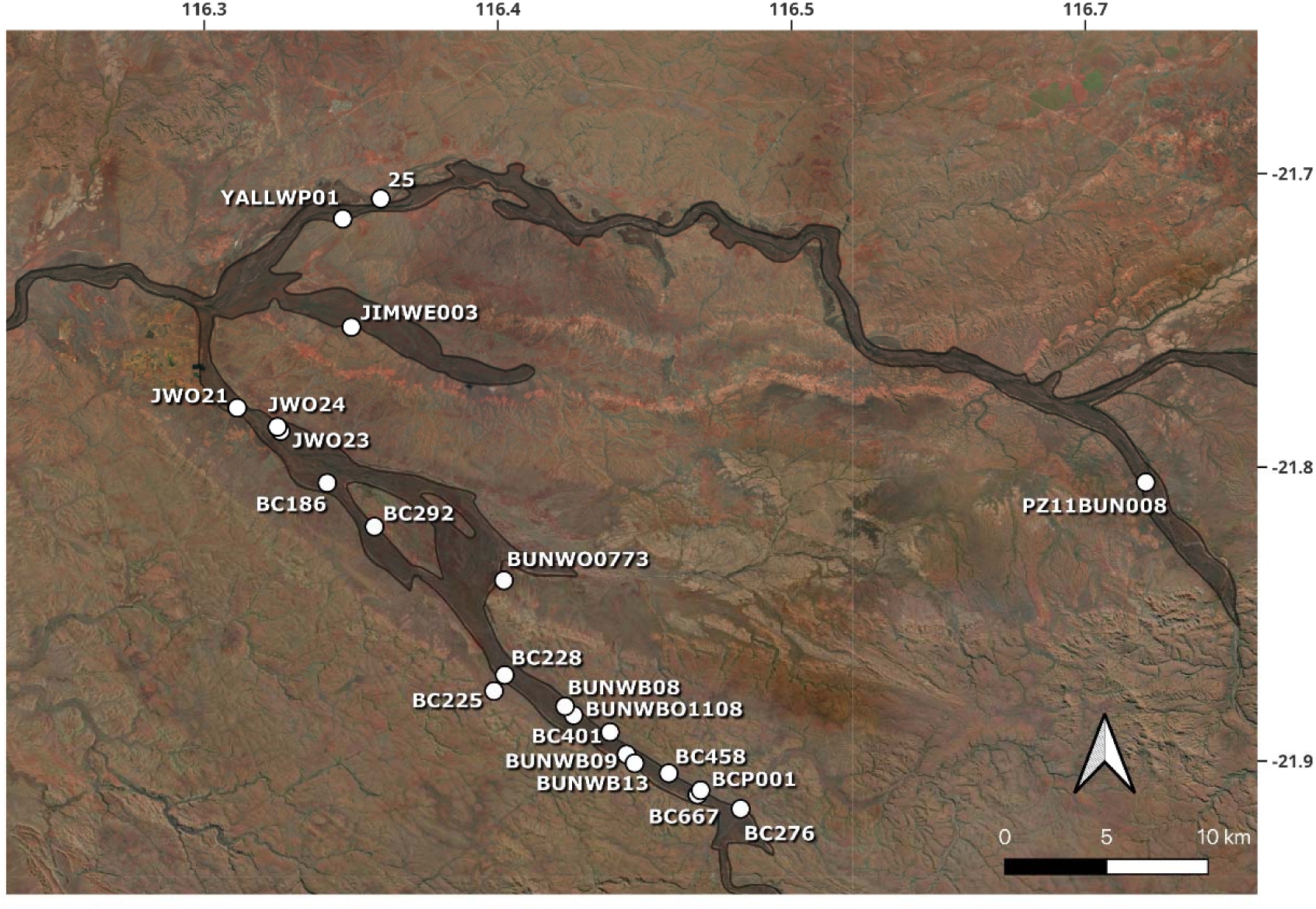
Map showing locations of the boreholes at the Bungaroo Creek borefield site, sampled for eDNA from groundwater, with bore locations and names. Map created using ArcMap v 10.3.1 (ESRI 2017), data layers provided by Rio Tinto.

Whole specimens were also collected at Bungaroo Creek by Guzik *et al*. (2024). These specimens were sequenced with long strand sequencing to confirm lineages present relative to the species identification with morphology. Sequences were used as an independent custom database to compare and test the efficacy of the GenBank records and the BRL developed here. These sequences are now available as part of the subterranean BRL (dx.doi.org/10.5883/DS-SUBFAUNA).

### eDNA extraction, assay, and bioinformatics

Entire membranes were extracted to ensure eDNA stability for stored samples. DNA was extracted from the water filters of samples and extraction/lab controls (i.e., extractions that use all reagents but no template DNA) using a DNeasy Blood and Tissue kit (Qiagen; Venlo, Netherlands) with the following modifications to the manufacturer’s protocol. The volume of tissue lysis buffer and proteinase K were doubled to ensure the entire filter membrane was adequately exposed to lysis during the 12 h incubation. This modification was conducted prior to loading the membrane digests into an automated QIAcube (Qiagen; Venlo, Netherlands) DNA extraction system. The eDNA extracts were eluted in 100 µl AE buffer. Four metabarcoding assays were applied to samples: nuclear *18S* rRNA (18S), and mitochondrial *16S* rRNA (Ins16S), *16S* rRNA (Crust16S) and *COI* (fwh) (see Table 1 for respective references). The four metabarcoding assays were not applied to all samples as some were used to test various assays; details on the samples sequenced using each assay can be found in (Supplementary Material 3). In general, the *18S* assay was applied to most of the samples in 2017 (*n* = 64) and 2021 (*n* = 67), *16S* rRNA to a subset of the 2017 samples (Crust 16S *n* = 37; Ins 16S *n* = 35), and *COI* to a subset of the 2021 samples (*n* = 16). Quantitative PCR (qPCR Applied Biosystems, USA) was then used to verify the quantity and quality of eDNA. PCR-amplifications were in 25 μl volumes containing: 1X AmpliTaq Gold®PCR buffer (Life Technologies, Massachusetts, USA), 2.5 mM MgCl2, 0.25 mM dNTPs, 0.4 μM each of forward and reverse primers (Integrated DNA Technologies, Australia), 0.4 mg/mL BSA (Fisher Biotec, Australia), 0.6 μl of 5X SYBR® Green (Life Technologies), 1 U AmpliTaq Gold® DNA Polymerase (Life Technologies), 2 μl of genomic DNA template, and made to volume with Ultrapure Distilled Water (Life Technologies). The cycling conditions were initial denaturation at 95° C for 5 min, followed by 40-50 cycles of 95° C for 30 s, annealing at primer specific temperature for 30 s (*16S* 52°C, *18S* 52°C, *COI* 51°C), 72° C for 45 s, and a final extension at 72° C for 10 min.

**Table 1:**
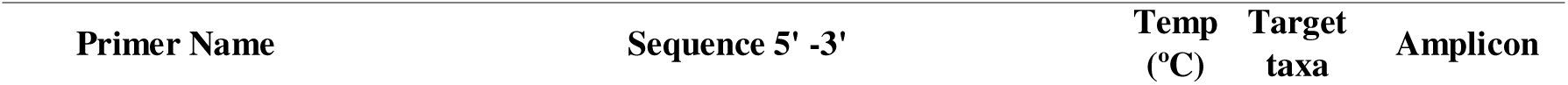
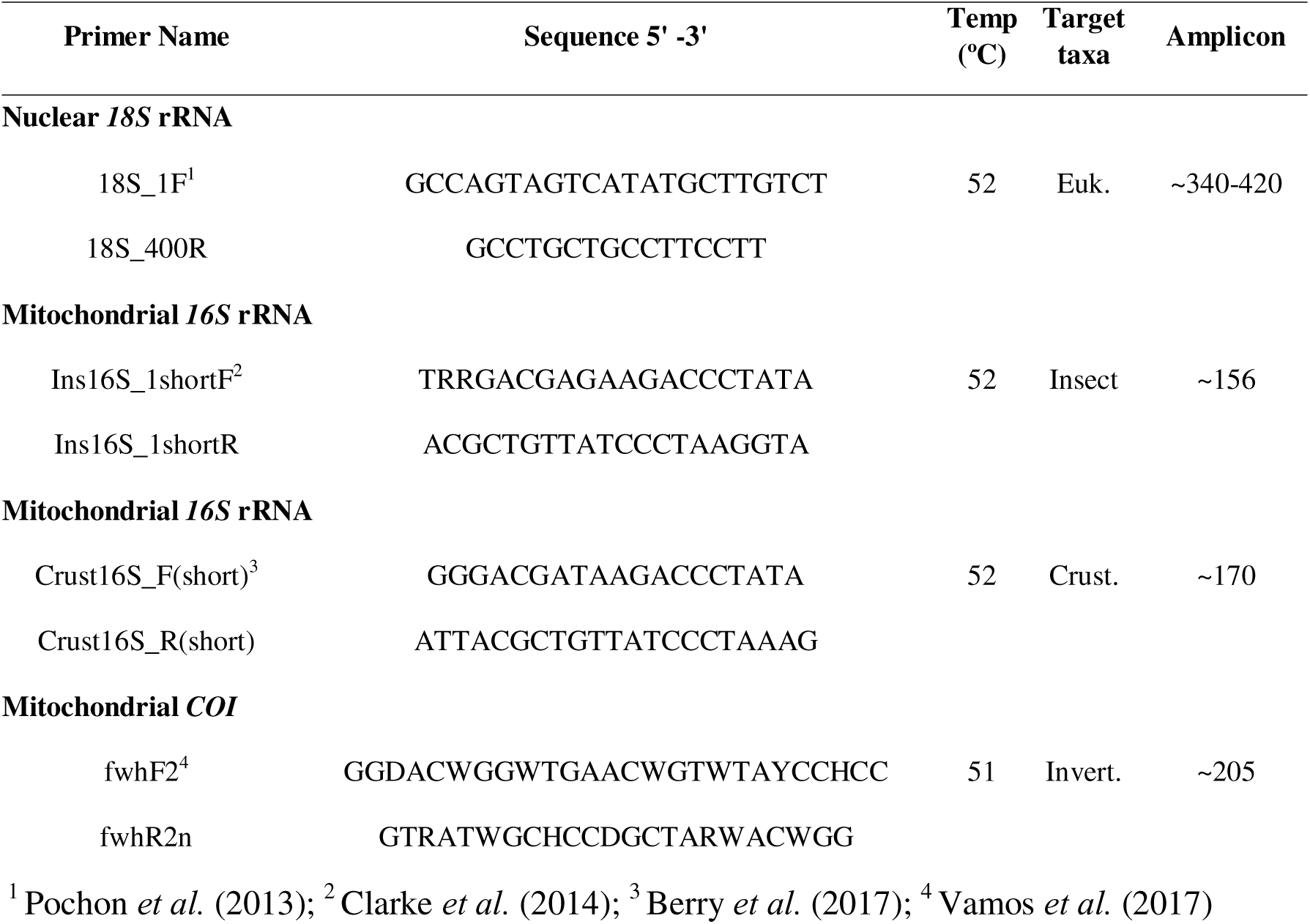
Assay primers and amplification details used in the present study specific to certain taxonomic groups and barcode genes, namely nuclear *18S* rRNA (eukaryotes^1^), mitochondrial *16S* rRNA (insects^2^ and crustaceans^3^) and mitochondrial *COI* (invertebrates^4^).

Samples were assigned a unique 6–8-base pair (bp) multiplex identifier tag (MID tag) and fusion PCR was used with the same conditions as described above to generate amplicons with sequencing primers and MID tags. MID-tag PCR amplicons were generated in duplicate, pooled together, quantified using a Qubit fluorometer (Thermo Fisher Scientific, Australia), and combined in equimolar concentrations. The libraries were then size selected using a PippinPrep (Millennium Science Pty Ltd., Australia) set at 160–500 bp (*16S* library), 200-500 bp (*COI* library) and 250–600 bp (*18S* library) and sequenced on a MiSeq following standard Illumina protocols. Bioinformatic analyses were performed on the Pawsey Supercomputer (Perth, WA; see Acknowledgements) using eDNAFlow, an automated workflow for the analysis of eDNA metabarcoding sequences (Mousavi-Derazmahalleh *et al*. 2021). Briefly, this workflow demultiplexes and filters sequences using the OBITools software package (Boyer *et al*. 2016) and USEARCH (Edgar 2016) for denoising to generate zero-radius operational taxonomic units (ZOTUs). A ZOTU is a cluster of DNA sequences that are very similar and can be seen as a rough proxy of a species (Mächler *et al*. 2021a; Mächler *et al*. 2021b). For the *COI* assay, sequences were clustered using USEARCH *cluster_otus* into OTUs with 97% sequence similarity due to the variability of the gene compared to ribosomal markers assayed here.

### Custom BLAST for Pilbara BRL and NCBI GenBank

Custom reference libraries for *18S*, *COI* and *16S* sequences were curated using the *refdb* package (Keck and Altermatt 2023) in R. We removed gaps in sequences, filtered by length (*CO1* min 400, max 900 bp; *18S* min 700, max 2000 bp; *16S* min 350, max 600 bp), harmonized taxonomy, and removed duplicate sequences. Further curation involved checking all taxonomic conflicts, including updating names for published synonyms (e.g., the schizomid *Paradraculoides* (Harvey *et al*. 2008) to *Draculoides* (Harvey 1992)), adding taxonomy for some sequences submitted with coarse identifications to GenBank or BOLD (e.g., Copepoda), and removing taxonomic identifications for sequences that were duplicated/represented by multiple individuals (e.g., *Draculoides* in *18S* sequences). In future, any downloads of the database need to repeat this editing. Details for all corrections can be found in supplementary files which include files for the current original and curated identifications (Supplementary Material 4). All ZOTUs were searched against the custom BRL database and GenBank (Benson *et al*. 2005) separately and together using BLASTn via the Pawsey Supercomputer. Search results were limited to 85% of bases being identical to the reference, ignoring alignments below query coverage of 95% and with 10 target sequences. For *18S* sequences, the first 50 bp of ZOTUs were removed to ensure the ZOTU sequences were consistent with the regions sequenced for the custom BRL. The importance of trimming these bases is that percent coverage is reduced when bases are included, increasing the mismatch between the ZOTU BLAST and database. This was especially important for *18S*. Taxonomic assignment was performed using lowest common ancestor (LCA) algorithm with eDNAFlow (Mousavi-Derazmahalleh *et al*. 2021). The parameters for this assignment were minimum 99% query coverage and 95% identity.

## Results

### BOLD – GenBank library

We curated a total of 5910 sequences from GenBank (*n*=4733) from 55 families of subterranean taxa. We captured GenBank and BOLD sequences for six barcode genes for *COI*-5P (i.e. 5’) (*n* = 4454), *16S* (*n* = 202), *18S* (*n* = 595), *28S* (*n* = 466), *12S* (*n* = 137), *COI*-3P (*n* = 56) for a total of 1462 BINs). The phylogeny for *COI*-5P (*n* = 4454) sequences was estimated using a Neighbor Joining approach (Figure 3). Nucleotide composition and saturation of first and third codon positions have not been accounted for in phylogenetic inference. The final reference libraries post-*refdb* curation included 116 sequences for *16S*, 249 sequences for *18S*, and 3139 sequences for *COI* (Supplementary Material 4).

**Figure 3:**
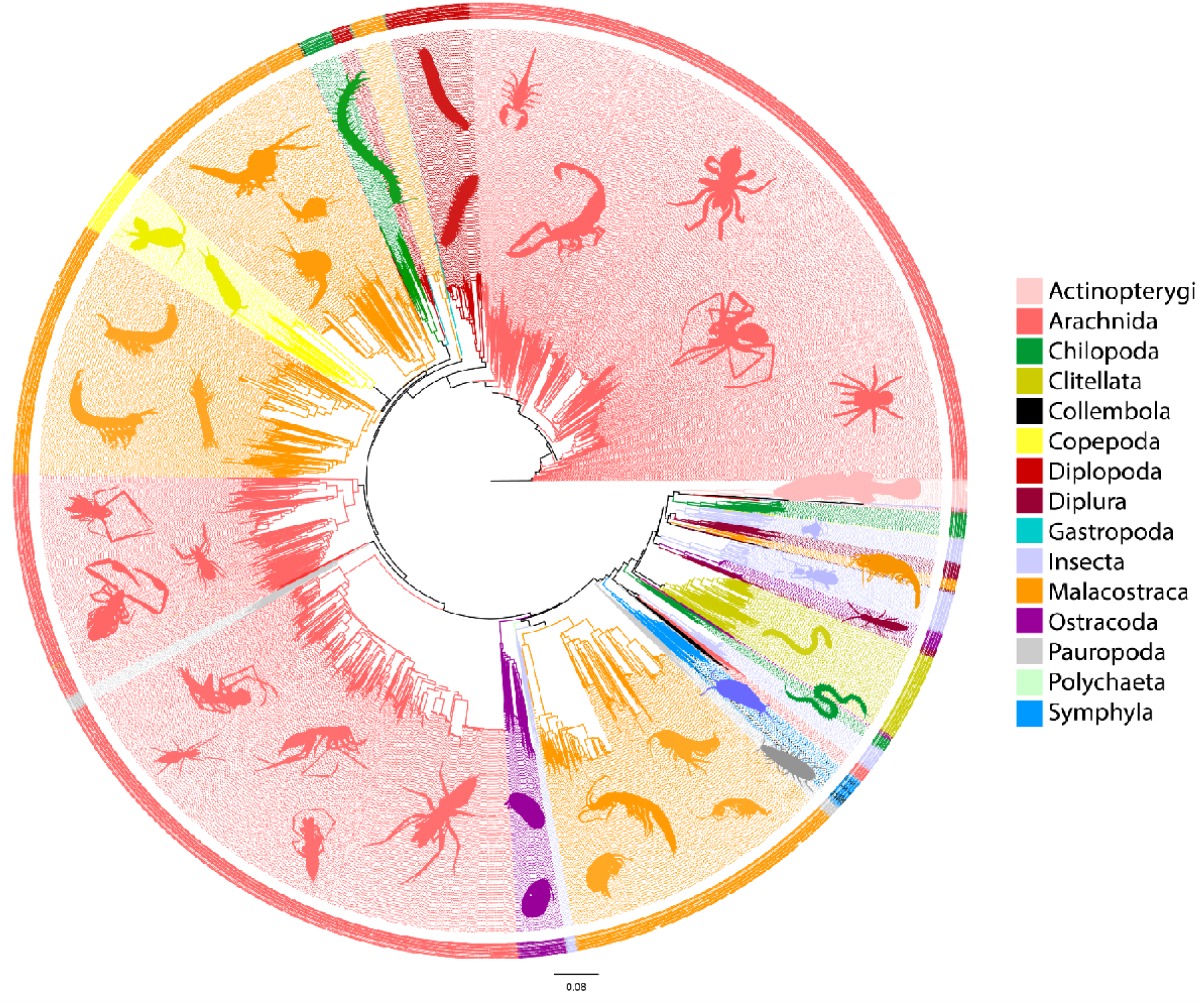
Pilbara subterranean fauna Custom Barcode Reference Library. Phylogenetic tree reconstructed from 5910 *COI* sequences of subterranean fauna curated from BOLD and GenBank using the *Taxon ID Tree* function to apply the neighbour-joining algorithm (Saitou and Nei 1987) under a Kimura 2-parameter model (Kimura 1980) in BOLD. Caution must be exercised when evaluating this tree, remembering it is a visual representation of the data, rather than accurate phylogenetic relationships.

### eDNA results

The *COI* assay generated 1,086,652 filtered reads from 16 samples corresponding to 718 ZOTUs clustered at 97% identity (Table 3). Similar to the *16S* results, only 12% of *COI* ZOTUs were identified using GenBank, 4.5% to fungi, 3.6% to arthropods, and the rest to miscellaneous eukaryotes and bacteria (Supplementary Material 5). Using the custom BRL, this taxonomic resolution increased such that six copepod ZOTUs, three amphipod ZOTUs, and a single snail ZOTU (Littorinimorpha) were identified (Supplementary Material 6-7). Several ZOTUs were identified using both the BRL and GenBank, but our BRL again provided greater taxonomic resolution. For example, a ZOTU identified to family level using GenBank could be resolved to a specific BOLD BIN to genus level when our BRL was implemented. However, numerous taxonomic groups were not detected with all markers and databases (Figure 3; Supplementary Material 8), namely many crustacean taxa.

**Table 3:** Total number of ZOTUs for eDNA groundwater samples collected at Bungaroo Creek, Pilbara, Western Australia for three gene regions and the number of successful hits of ZOTU’s to a. metazoa and b. subterranean fauna for each database (**GenBank**; our custom **BRL**; **both** databases combined, and taxa that were **shared** between the two databases).

|  |  | 18S | 16S | COI |
| --- | --- | --- | --- | --- |
| **a. Total metazoa** | ZOTUs | 4024 | 474 | 718 |
|  | GenBank | 2883 | 42 | 81 |
|  | BRL | 10 | 17 | 10 |
|  | both | 2885 | 58 | 87 |
|  | shared | 8 | 0 | 4 |
| **b. Subterranean fauna** | GenBank | 112 | 0 | 6 |
|  | BRL | 10 | 17 | 10 |
|  | both | 117 | 17 | 12 |
|  | shared | 3 | 0 | 4 |

In the *18S* dataset, we generated 4,540,472 filtered reads corresponding to 4024 ZOTUs from 131 samples (Table 3). Of the 2885 ZOTUs with hits above 95% identity, 35% were assigned to fungi, 13.7% to ciliates, 7.4% to arthropods, 5.5% to streptophyte plants, and 15% to the broader Eukaryota by the LCA taxonomic assignment algorithm (Supplementary Material 5). The ZOTUs identified using the BRL were primarily arachnids (4 ZOTUs to Schizomida), bathynellids (4 ZOTUs), with a single ZOTU each identified as a micro shrimp (Thermosbaenacea) and snail (Littorinimorpha) (Supplementary Material 9-10) and (Figure 4; Supplementary Material 8).

**Figure 4:**
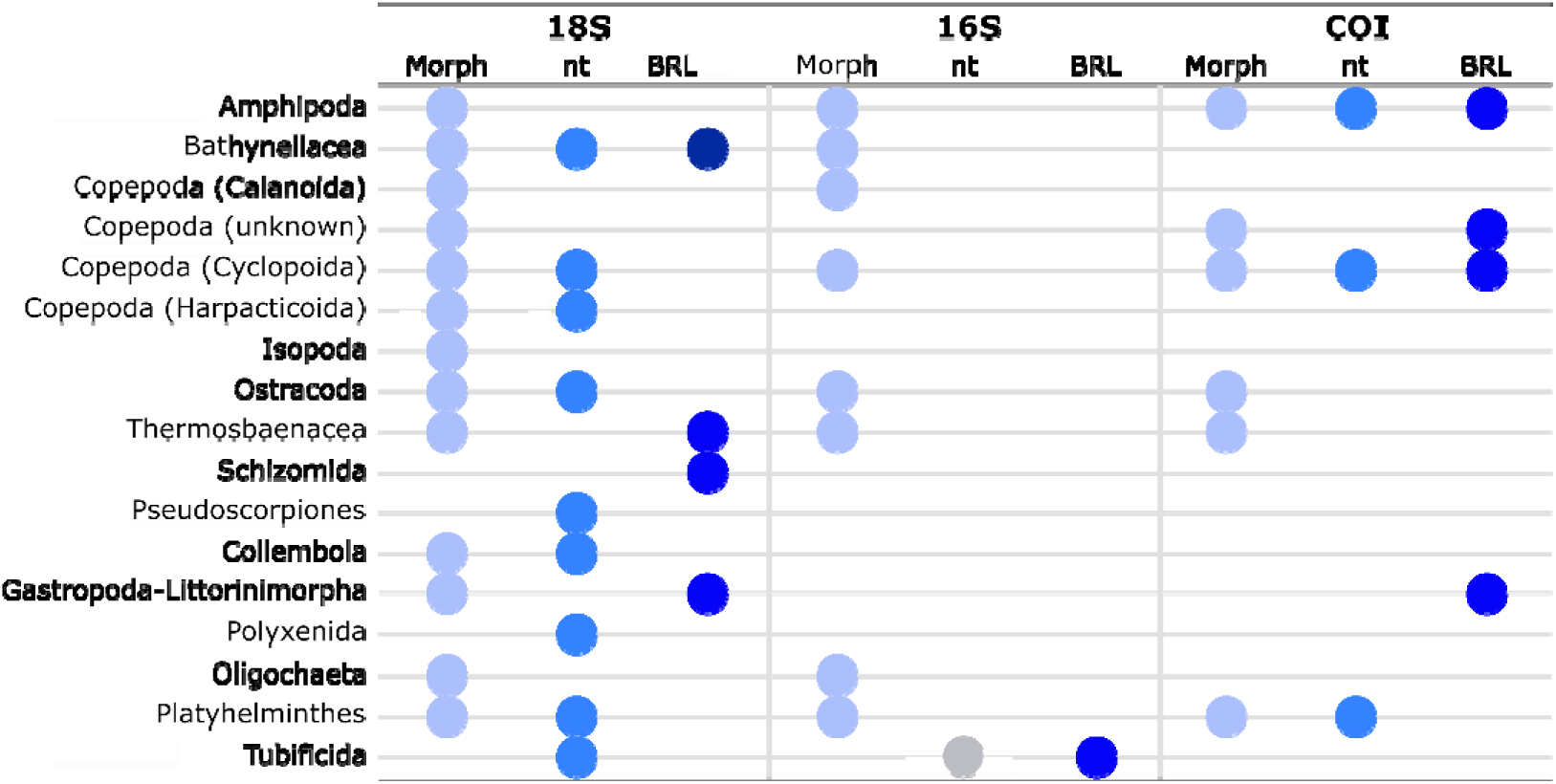
Presence or absence of detection of subterranean orders from eDNA (*medium and dark blue*) and morphology (*light blue*) where circles represent subterranean orders detected using eDNA metabarcoding in that gene region, while no circle represents taxa not detected. The *grey circle* indicates that a taxon in the same order or class was detected using eDNA. nt: nucleotide GenBank BLAST search; BRL: subterranean custom barcode reference library.

We generated 378,069 filtered reads for the *16S* eDNA data with 474 ZOTUs from 37 samples using the crustacean and insect assays (Table 3). Of these, 58 ZOTUs (12%) were assigned taxonomic information (Supplementary Material 5). The *16S* ZOTUs identified using **GenBank** were almost all (34/42) assigned to Insecta (Supplementary Material 5), but these were all identified as surface terrestrial taxa. In contrast, using the custom **BRL** developed here, we were able to identify 17 *16S* ZOTUs as subterranean annelid worms (Clitellata and Tubificida) (likely due to the assay primer redundancy) (Supplementary Material 11-12) (Figure 3; Supplementary Material 8).

## Discussion

Here we present the first curated sequence database for the identification of subterranean fauna from eDNA, comprising 5910 sequences from 39 invertebrate orders (Figure 3). DNA sequence data and associated genetic diversity can be used as molecular proxies for biodiversity (Floyd *et al*. 2002; Blaxter and Floyd 2003) and can be especially important for identification of cryptic and unknown species (King *et al*. 2012; Murphy *et al*. 2015; King *et al*. 2022). This is even more important for subterranean realms, which at times receive substantial conservation attention in Australia due to interconnected industry interest and legislative requirements (Gibson *et al*. 2019; Saccò *et al*. 2022a). A custom BRL requires DNA sequences at both broad (order, family) and narrow (genus to population level) taxonomic coverage to be useful in identifying unknown individuals and sequences in genetic and taxonomic studies (Hebert *et al*. 2013). All sequences were curated in a BOLD project, which will, from here on, form the basis of a FAIR publicly available eDNA resource that can be used to reference groundwater and soil eDNA metabarcoding OTUs. We also tested our BRL with eDNA metabarcoding OTUs and demonstrated its great potential for improving identification and taxonomic resolution of anonymous OTUs. Given the extraordinary biodiversity value of subterranean fauna and ongoing human impacts on groundwater-dependent ecosystems (Saccò *et al*. 2024), a coordinated effort is required to advance such data infrastructure and aid future conservation management planning. Our view is that new data can be incorporated into the subterranean BRL by researchers over time, thereby creating a dynamic and expandable platform for a variety of stakeholders that can accommodate diverse metadata. This database forms a framework for new and future sequence entries and represents the first step towards a comprehensive publicly accessible, global reference library for subterranean fauna.

### Towards a subterranean BRL: curation of existing data from GenBank and BOLD

In curating the custom database presented here, we made two key observations with respect to existing and published data. First, *COI* dominated the subterranean fauna legacy data in both GenBank and BOLD (i.e., curated data: *COI*-5P (4454), *16S* (202), *18S* (595), *28S* (466), *12S* (137), *COI*-3P (56)). Historically, *COI* has been heavily used for subterranean fauna due to accessibility of this gene using standard universal invertebrate primers (Folmer *et al*. 1994; Simon *et al*. 1994) and its usefulness for discriminating at the species level to delineate new and undescribed species (Hebert *et al*. 2003). Furthermore, in WA specifically, the WA Environmental Protection Authority’s technical guidance for subterranean fauna surveys for environmental impact assessment guidance (Environmental Protection Authority 2021) has led to *COI* being frequently implemented for molecular species identification and verification of unknown species, resulting in a large number of these sequences being deposited in GenBank (Biologic Environmental Survey 2021). Second, taxonomic coverage in the BRL was also uneven for arthropods with arachnids and crustaceans being heavily represented, likely due to a bias in taxonomic expertise among researchers and funding availability (e.g. Jarman and Elliot 2000; Leys *et al*. 2003; Cooper *et al*. 2007; Finston *et al*. 2007; Page *et al*. 2007; Cooper *et al*. 2008; Guzik *et al*. 2008; Leys and Watts 2008; Page *et al*. 2008; Guzik *et al*. 2009; Bradford *et al*. 2010; Chakrabarty 2010; Abrams *et al*. 2012; Asmyhr and Cooper 2012; Cook *et al*. 2012; Karanovic and Cooper 2012; Murphy *et al*. 2012; Abrams *et al*. 2013; Larson *et al*. 2013; Murphy *et al*. 2013; Asmyhr *et al*. 2014; Javidkar *et al*. 2015; Karanovic *et al*. 2015; Camacho *et al*. 2018; Moore *et al*. 2018; Matthews *et al*. 2020) and as the most abundant groups in these systems. Overall, the BRL contains the most comprehensive DNA sequence database for subterranean fauna currently available from the Pilbara region. Nonetheless, we reiterate that much of this biota remains uncharacterised globally, further highlighting a need for the development of a BRL for the identification of subterranean fauna worldwide.

### BOLD as a DNA database platform

To facilitate curation of a subterranean fauna database, we took advantage of the existing database platform, iBOL’s BOLD. Using this infrastructure has provided several advantages over the establishment of a new database, including: 1) investment of energy and resources into establishing a new database is rarely feasible or practical; 2) BOLD is fully funded and maintained into the future, ensuring the resource developed here will be accessible and available to researchers in perpetuity; 3) the BRL curated in BOLD addresses almost all the FAIR principles of (Wilkinson *et al*. 2016; Stall *et al*. 2019) required for curating informative and accessible online resources; and 4) BOLD allows the submission of multiple barcoding genes such as *16S* and *18S* for comparison against other eDNA metabarcoding assays. In particular, this applies to the functionality of BOLD as a user-friendly web interface in which both public and private projects and databases can be created and where taxonomic, geographic, and photographic metadata can be associated with individual sequences (Ratnasingham and Hebert 2007). While other repositories, such as GenBank, provide several of these features, most are not associated with sequence data once it is downloaded. We also note BOLD supports GenBank data formats and allows visualisation of associated metadata for the sample in question. Finally, BOLD hosts an *in silico* delimitation algorithm that assigns identifiers (BINs) to *COI* lineages based on the genetic distances between all sequences in the database. BINs can provide a centralised system for identifying lineages equivalent to species (Hajibabaei *et al*. 2007). While BINs may be unavailable for underrepresented taxonomic groups and geographic regions (Lahaye *et al*. 2008) or specific genes such as *16S* and can change over time, we consider the metric to be an important tool for unifying nomenclature of undescribed species (Guzik *et al*. 2024).

### Validation of the reference library with eDNA metabarcoding ZOTUs

Each sequence cluster (i.e., ZOTU) estimated from eDNA metabarcoding is represented by thousands of reads depending on the ubiquity of organisms in the environment (Takahara *et al*. 2012), shedding rate (Andruszkiewicz Allan *et al*. 2021), and longevity of eDNA (Ruppert *et al*. 2019) for each organism. However, the ZOTUs generated from eDNA metabarcoding samples are anonymous upon sequencing. In order to effectively identify species from ZOTUs, a well sampled BRL is required. Here, for the first time, we tested a new subterranean BRL with eDNA metabarcoding ZOTUs. We observed that using the custom BRL improved taxonomic identifications for all assays tested here. The custom BRL allowed identification of *COI* OTUs to lower taxonomic rank (i.e. genus and BOLD BIN rather than family). In all three barcoding regions, the custom BRL identified subterranean ZOTUs that could not be identified from GenBank alone (Table 3). However, only a small proportion (12%) of *16S* and *COI* OTUs were able to be identified by either database compared to *18S* (72%), perhaps a result of increased molecular discrimination between barcodes in those genes (*16S* and *COI*). In the *18S* blast results, it was common to get hits with >97% identification for different phyla/orders, which then have to be dropped to the lowest common ancestor. Some of these results may also arise from poor database curation as well as limited molecular discrimination since most individuals/sequences are not identified to species or are incorrectly identified (Ruppert *et al*. 2019). It was observed that major taxonomic gaps in the eDNA data still exist (Figure 3). Additionally, the curated custom BRL only included 116 reference sequences for *16S*. We can expect greater rates of identification as this database grows. Together the BRL and GenBank were successful in characterising most of the information (Guzik *et al*. 2024).

A major limiting factor of BOLD for hosting the subterranean BRL was in testing the applicability of the database by comparing BLAST queries of eDNA ZOTUs against the custom BRL. Currently, a BLAST of the custom database cannot be run from within BOLD, aside from the *BOLD Identification System (IDS)* for *COI*, which returns a species-level identification when one is possible. Instead we used the R package *refdb* to audit and manage the custom database (Keck and Altermatt 2023). In executing this package, we observed a number of taxonomic inconsistencies (e.g., taxonomic information that has not been updated from more recent revisions) that were manually amended. Whilst we provide the static BRL files in Supplementary Material 3, unfortunately, these edits will again need to be made following any downloads of future updates of the database by other users, unless the original authors of the sequence update their records and provide updated versions. It is expected that future development of BOLD will include opportunities to BLAST custom databases from within the platform as per the online platform mBRAVE (Ratnasingham 2019).

### A plan for future development of a global custom BRL for subterranean fauna

Using FAIR principles of data infrastructure, we aimed to establish an information resource for eDNA biomonitoring of subterranean fauna that is shared and dynamic. In particular, we aimed to inform a unified nomenclature for subterranean fauna and allow for stable and repeatable queries for taxonomic assignment of OTUs generated from eDNA metabarcoding data. The Pilbara BRL curated here showed that while there are still major gaps in taxonomy of subterranean fauna and available DNA sequence data in both BOLD and GenBank (Figure 3), there is great potential for improving identification of anonymous OTUs from eDNA data. Other studies have found similar results (King *et al*. 2022; Perina *et al*. 2023) and we suggest that further collaborative research should prioritise expanding and curating a global subterranean BRL that would undertake the following tasks to increase the functionality of the BRL for eDNA metabarcoding as these methods develop (see Figure 5):

1. Further curate sequence data from *existing* public databases, such as GenBank and BOLD, for subterranean fauna globally to establish the phylogenetic backbone for a custom reference library.
2. Promote submission of data by academic and non-academic stakeholders from unpublished surveys, reports and publications with the recognition that some vetting of data may be required.
3. Encourage collaborative taxonomic research on key groups and a global update of taxonomic nomenclature in existing public databases by original authors.
4. Identify key taxonomic groups missing from GenBank and BOLD for development of future DNA sequencing projects.
5. Identify key geographic regions missing from GenBank and BOLD for development of future DNA sequencing projects.
6. Facilitate future targeted sequencing of morphologically identified specimens for target groups using either whole mitochondrial genome sequencing/skimming or long read sequencing of a recommended panel of loci (e.g. *18S*, *COI*, *16S* and *12S*).

**Figure 5:**
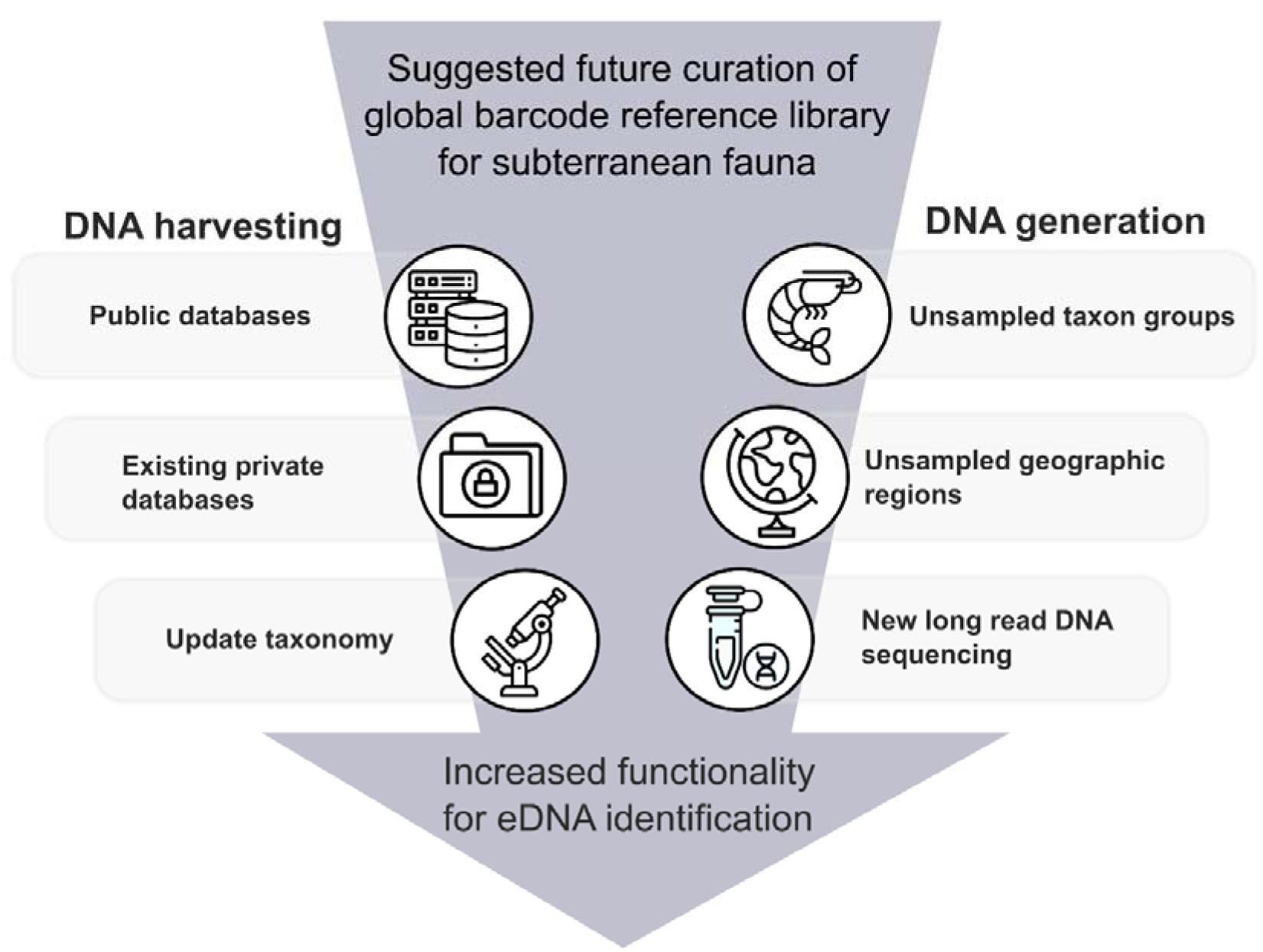
A framework for expanding and curating a global subterranean BRL for improved reference functionality in eDNA OTU identification.

We indicate that the BOLD database is a suitable data repository because of its open access, updatable format, metadata framework and ability to formulate a systematic framework for fauna (see also Guzik *et al*. 2024). Further, in the absence of formal taxonomy for many subterranean species we have proposed that the assignment of BINs in BOLD may be a solution for a unified nomenclature for subterranean fauna (Guzik *et al*. 2024).

### Conclusions

The implementation of high throughput eDNA metabarcoding for biodiversity assessment and monitoring is of intense interest for its augmentation of traditional monitoring and survey methods (Thomsen *et al*. 2012; Bohmann *et al*. 2014; Deiner *et al*. 2016; Goldberg *et al*. 2016). This is especially true for monitoring endangered, rare, and elusive taxa (Bohmann *et al*. 2014; Furlan *et al*. 2016), including subterranean fauna (White *et al*. 2020). A robust BRL is essential for effective translation of OTUs to taxonomic meaning for the detection of fauna using metabarcoding methods. Here we present a major step forward in establishing a BRL which we hope will revolutionise subterranean faunal community assessment, monitoring and species identification. We recognise that much data is not available on public reference databases and we hope this project will foster timely new initiatives to facilitate the development of a global BRL for subterranean fauna.

## Acknowledgements

This research was part funded by Australian Research Council (ARC) Linkage Project Grant LP140100555 to A.D.A. and S.J.B.C. (linkage partners: South Australian Museum, Western Australian Museum, Department of Biodiversity, Conservation and Attractions (formerly Department of Parks and Wildlife), Bennelongia Pty Ltd, Biota Environmental Sciences Pty Ltd), ARC Linkage Project Grant LP190100555 to A.D.A. and S.J.B.C. (linkage partners: Rio Tinto, BHP, Chevron, South Australian Museum, Western Australian Museum, Department of Biodiversity, Conservation and Attractions, Department for Water and Environmental Regulation, Western Australian Biodiversity Science Institute (WABSI)), Rio Tinto consulting contract to Nicole White and Mike Bunce, and a research contract with WABSI funded by Rio Tinto and BHP. This work was supported by resources provided by the Pawsey Supercomputing Centre with funding from the Australian Government and the Government of Western Australia.

## References

1. Abrams K., Guzik M., Cooper S., Humphreys W., King R., Cho J.-L., Austin A. (2012) What lies beneath: molecular phylogenetics and ancestral state reconstruction of the ancient subterranean Australian Parabathynellidae (Syncarida, Crustacea). Molecular Phylogenetics and Evolution 64(1), 130–144.

2. Abrams K. M., King R. A., Guzik M. T., Cooper S. J. B., Austin A. D. (2013) Molecular phylogenetic, morphological and biogeographic evidence for a new genus of parabathynellid crustaceans (Syncarida: Bathynellacea) from groundwater in an ancient southern Australian landscape. Invertebrate Systematics 27(2), 146–172.

3. Aizpurua O., Budinski I., Georgiakakis P., Gopalakrishnan S., Ibanez C., Mata V., Rebelo H., Russo D., Szodoray-Paradi F., Zhelyazkova V., Zrncic V., Gilbert M. T. P., Alberdi A. (2018) Agriculture shapes the trophic niche of a bat preying on multiple pest arthropods across Europe: Evidence from DNA metabarcoding. Molecular Ecology 27(3), 815–825.

4. Andruszkiewicz Allan E., Zhang W. G., C Lavery A., F Govindarajan A. (2021) Environmental DNA shedding and decay rates from diverse animal forms and thermal regimes. Environmental DNA 3(2), 492–514.

5. Asmyhr M. G., Cooper S. J. B. (2012) Difficulties barcoding in the dark: the case of crustacean stygofauna from eastern Australia. Invertebrate Systematics 26(6), 583–591.

6. Asmyhr M. G., Linke S., Hose G., Nipperess D. A. (2014) Systematic conservation planning for groundwater ecosystems using phylogenetic diversity. PLOS ONE 9(12), e115132.

7. Belle C. C., Stoeckle B. C., Geist J. (2019) Taxonomic and geographical representation of freshwater environmental DNA research in aquatic conservation. Aquatic Conservation: Marine and Freshwater Ecosystems 29(11), 1996–2009.

8. BejaCPereira, A. L. B. A. N. O., Oliveira, R., Alves, P. C., Schwartz, M. K., and Luikart, G. (2009). Advancing ecological understandings through technological transformations in noninvasive genetics. Molecular Ecology Resources 9(5), 1279–1301.

9. Beng K. C., Corlett R. T. (2020) Applications of environmental DNA (eDNA) in ecology and conservation: opportunities, challenges and prospects. Biodiversity and Conservation 29, 2089–2121.

10. Berry O., Jarman S., Bissett A., Hope M., Paeper C., Bessey C., Schwartz M. K., Hale J., Bunce M. (2021) Making environmental DNA (eDNA) biodiversity records globally accessible. Environmental DNA 3(4), 699–705.

11. Berry T. E., Osterrieder S. K., Murray D. C., Coghlan M. L., Richardson A. J., Grealy A. K., Stat M., Bejder L., Bunce M. (2017) DNA metabarcoding for diet analysis and biodiversity: A case study using the endangered Australian sea lion (Neophoca cinerea). Ecology and Evolution 7(14), 5435–5453.

12. Biologic Enivronmental Survey (2021) McPhee Creek Subterranean Fauna Assessment.

13. Bissett A., Fitzgerald A., Court L., Meintjes T., Mele P. M., Reith F., Dennis P. G., Breed M. F., Brown B., Brown M. V., Brugger J., Byrne M., Caddy-Retalic S., Carmody B., Coates D. J., Correa C., Ferrari B. C., Gupta V. V., Hamonts K., Haslem A., Hugenholtz P., Karan M., Koval J., Lowe A. J., Macdonald S., McGrath L., Martin D., Morgan M., North K. I., Paungfoo-Lonhienne C., Pendall E., Phillips L., Pirzl R., Powell J. R., Ragan M. A., Schmidt S., Seymour N., Snape I., Stephen J. R., Stevens M., Tinning M., Williams K., Yeoh Y. K., Zammit C. M., Young A. (2016) Introducing BASE: the Biomes of Australian Soil Environments soil microbial diversity database. GigaScience 5, 21.

14. Bista I., Carvalho G. R., Walsh K., Seymour M., Hajibabaei M., Lallias D., Christmas M., Creer S. (2017) Annual time-series analysis of aqueous eDNA reveals ecologically relevant dynamics of lake ecosystem biodiversity. Nature Communications 8, 14087.

15. Blaxter M., Floyd R. (2003) Molecular taxonomics for biodiversity surveys: already a reality. Trends in Ecology and Evolution 18(6), 268–269.

16. Bohmann K., Evans A., Gilbert M. T., Carvalho G. R., Creer S., Knapp M., Yu D. W., de Bruyn M. (2014) Environmental DNA for wildlife biology and biodiversity monitoring. Trends Ecology and Evolution 29(6), 358–67.

17. Boulton A. J., Fenwick G. D., Hancock P. J., Harvey M. S. (2008) Biodiversity, functional roles and ecosystem services of groundwater invertebrates. Invertebrate Systematics 22(2), 103–116.

18. Boykin L. M., Armstrong K., Kubatko L., De Barro P. (2012) DNA barcoding invasive insects: database roadblocks. Invertebrate Systematics 26(6).

19. Bradford T., Adams M., Humphreys W. F., Austin A. D., Cooper S. J. (2010) DNA barcoding of stygofauna uncovers cryptic amphipod diversity in a calcrete aquifer in Western Australia’s arid zone. Molecular Ecology Resources 10(1), 41–50.

20. Bridge P. D., Roberts P. J., Spooner B. M., Panchal G. (2003) On the unreliability of published DNA sequences. New Phytologist 160(1), 43–48.

21. Butchart S. H., Walpole M., Collen B., Van Strien A., Scharlemann J. P., Almond R. E., Baillie J. E., Bomhard B., Brown C., Bruno J., Carpenter K. E. (2010) Global biodiversity: Indicators of recent declines. Science 328(5982), 1164–1168.

22. Camacho A. I., Mas-Peinado P., Dorda B. A., Casado A., Brancelj A., Knight L. R., Hutchins B., Bou C., Perina G., Rey I. (2018) Molecular tools unveil an underestimated diversity in a stygofauna family: a preliminary world phylogeny and an updated morphology of Bathynellidae (Crustacea: Bathynellacea). Zoological Journal of the Linnean Society 183(1), 70–96.

23. Chakrabarty P. (2010) Status and phylogeny of Milyeringidae (Teleostei: Gobiiformes), with the description of a new blind cave-fish from Australia, Milyeringa brooksi, n. sp. Zootaxa 2557(1), 19–28.

24. Clarke L. J., Soubrier J., Weyrich L. S., Cooper A. (2014) Environmental metabarcodes for insects: in silico PCR reveals potential for taxonomic bias. Molecular Ecology Resources 14(6), 1160–70.

25. Coissac E., Riaz T., Puillandre N. (2012) Bioinformatic challenges for DNA metabarcoding of plants and animals. Molecular Ecology 21(8), 1834–47.

26. Cook B., Abrams K., Marshall J., Perna C., Choy S., Guzik M., Cooper S. (2012) Species diversity and genetic differentiation of stygofauna (Syncarida: Bathynellacea) across an alluvial aquifer in north-eastern Australia. Australian Journal of Zoology 60(3), 152–158.

27. Cooper S. J. B., Bradbury J. H., Saint K. M., Leys R., Austin A. D., Humphreys W. F. (2007) Subterranean archipelago in the Australian arid zone: mitochondrial DNA phylogeography of amphipods from central Western Australia. Molecular Ecology 16, 1533–1544.

28. Cooper S. J. B., Saint K. M., Taiti S., Austin A. D., Humphreys W. F. (2008) Subterranean archipelago: mitochondrial DNA phylogeography of stygobitic isopods (Oniscidea: *Haloniscus*) from the Yilgarn region of Western Australia. Invertebrate Systematics 122, 195–203.

29. Couton M., Hürlemann S., Studer A., Alther R., Altermatt F. (2023a) Groundwater environmental DNA metabarcoding reveals hidden diversity and reflects landCuse and geology. Molecular Ecology 32(13), 3497–3512.

30. Couton M., Studer A., Hürlemann S., Locher N., Knüsel M., Alther R., Altermatt F. (2023b) Integrating citizen science and environmental DNA metabarcoding to study biodiversity of groundwater amphipods in Switzerland. Scientific Reports 13(1), 18097.

31. Creer S., Deiner K., Frey S., Porazinska D., Taberlet P., Thomas W. K., Potter C., Bik H. M., Freckleton R. (2016) The ecologist’s field guide to sequence-based identification of biodiversity. Methods in Ecology and Evolution 7(9), 1008–1018.

32. Cristescu M. E. (2014) From barcoding single individuals to metabarcoding biological communities: towards an integrative approach to the study of global biodiversity. Trends in Ecology and Evolution 29(10), 566–71.

33. Curry C. J., Gibson J. F., Shokralla S., Hajibabaei M., Baird D. J. (2018) Identifying North American freshwater invertebrates using DNA barcodes: are existing COI sequence libraries fit for purpose? Freshwater Science 37(1), 178–189.

34. Danielopol D. L., Griebler C., Gunatilaka A., Notenboom J. (2003) Present state and future prospects for groundwater ecosystems. Environmental Conservation 30(2), 104–130.

35. Deagle B. E., Jarman S. N., Coissac E., Pompanon F., Taberlet P. (2014) DNA metabarcoding and the cytochrome c oxidase subunit I marker: not a perfect match. Biology Letters 10(9).

36. Deiner K., Fronhofer E. A., Machler E., Walser J. C., Altermatt F. (2016) Environmental DNA reveals that rivers are conveyer belts of biodiversity information. Nature Communication 7, 12544.

37. Deharveng, L., Le, C.K., Bedos, A., Judson, M.L., Le, C.M., Lukić, M., Luu, H.T., Ly, N.S., Nguyen, T.Q.T., Truong, Q.T. and Vermeulen, J. (2023) A hotspot of subterranean biodiversity on the brink: Mo So Cave and the Hon Chong Karst of Vietnam. Diversity, 15(10), 1058.

38. Department of Climate Change E, the Environment and Water (1999) Environment Protection and Biodiversity Conservation Act.

39. DeWaard J. R., Ratnasingham S., Zakharov E. V., Borisenko A. V., Steinke D., Telfer A. C., Perez K. H., Sones J. E., Young M. R., Levesque-Beaudin V. (2019) A reference library for Canadian invertebrates with 1.5 million barcodes, voucher specimens, and DNA samples. Scientific Data 6(1), 308.

40. Dormontt E. E., Van Dijk K.-J., Bell K. L., Biffin E., Breed M. F., Byrne M., Caddy-Retalic S., Encinas-Viso F., Nevill P. G., Shapcott A. (2018) Advancing DNA barcoding and metabarcoding applications for plants requires systematic analysis of herbarium collections— an Australian perspective. Frontiers in Ecology and Evolution 6, 134.

41. Eberhard S. M., Halse S. A., Humphreys W. F. (2005) Stygofauna in the Pilbara region, north-west Western Australia: a review. Journal of the Royal Society of Western Australia 88, 167–176.

42. Edgar R. C. (2016) UNOISE2: improved error-correction for Illumina 16S and ITS amplicon sequencing. BioRxiv, 081257.

43. Environmental Protection Authority (2021) Technical guidance – Subterranean fauna surveys for environmental impact assessment. Depeartment of Environment and Water Regulation). (EPA: Western Australia).

44. Ferreira, R.L., Giribet, G., Du Preez, G., Ventouras, O., Janion, C. and Silva, M.S. (2020) The Wynberg cave system, the most important site for cave fauna in South Africa at risk. Subterranean Biology 36, 73–81.

45. Ficetola G. F., Miaud C., Pompanon F., Taberlet P. (2008) Species detection using environmental DNA from water samples. Biology Letters 4(4), 423–5.

46. Finston T. L., Johnson M. S., Humphreys W. F., Eberhard S. M., Halse S. A. (2007) Cryptic speciation in two widespread subterranean amphipod genera reflects historical drainage patterns in an ancient landscape. Molecular Ecology 16, 355–365.

47. Floyd R., Abebe E., Papert A., Blaxter M. (2002) Molecular barcodes for soil nematode identification. Molecular Ecology 11(4), 839–850.

48. Folmer O., Black M., Hoeh W., Lutz R., Vrijenoek R. (1994) DNA primers for amplification of mitochondrial cytochrome c oxidase subunit 1 from diverse metazoan invertebrates. Molecular Marine Biology and Biotechnology 3(5), 294–299.

49. Furlan E. M., Gleeson D., Hardy C. M., Duncan R. P. (2016) A framework for estimating the sensitivity of eDNA surveys. Molecular Ecology Resources 16(3), 641–54.

50. Gallão, J.E., Ribeiro, D.B., Gallo, J.S. and Bichuette, M.E., 2023. There and Back Again— The Igatu Hotspot Siliciclastic Caves: Expanding the data for subterranean fauna in Brazil, Chapada Diamantina Region. Diversity 15(9) 991.

51. Gibson L., Humphreys W. F., Harvey M., Hyder B., Winzer A. (2019) Shedding light on the hidden world of subterranean fauna: A transdisciplinary research approach. Science of The Total Environment.

52. Goldberg C. S., Turner C. R., Deiner K., Klymus K. E., Thomsen P. F., Murphy M. A., Spear S. F., McKee A., Oyler-McCance S. J., Cornman R. S., Laramie M. B., Mahon A. R., Lance R. F., Pilliod D. S., Strickler K. M., Waits L. P., Fremier A. K., Takahara T., Herder J. E., Taberlet P., Gilbert M. (2016) Critical considerations for the application of environmental DNA methods to detect aquatic species. Methods in Ecology and Evolution 7(11), 1299–1307.

53. Goricki S., Stankovic D., Aljancic M., Snoj A., Kuntner M., Gredar T., Vodnik L., Aljancic G. (2016) Searching for the black *Proteus* with the help of eDNA/Iskanje crnega mocerila s pomocjo okoljske DNA. Natura Sloveniae 18(1), 57.

54. Griebler C., Avramov M. (2015) Groundwater ecosystem services: a review. Freshwater Science 34(1), 355–367.

55. Guzik M. T., Abrams K. M., Cooper S. J. B., Humphreys W. F., Cho J.-L. (2008) Phylogeography of the ancient Parabathynellidae (Crustacea:Bathynellacea) from the Yilgarn region of Western Australia. Invertebrate Systematics 22, 205–216.

56. Guzik M. T., Austin A. D., Cooper S. J., Harvey M. S., Humphreys W. F., Bradford T., Eberhard S. M., King R. A., Leys R., Muirhead K. A., Tomlinson M. (2011a) Is the Australian subterranean fauna uniquely diverse? Invertebrate Systematics 24(5), 407–418.

57. Guzik M. T., Cooper S., Humphreys W. F., Austin A. D. (2009) Fine-scale comparative phylogeography of a sympatric sister species triplet of subterranean diving beetles from a single calcrete aquifer in Western Australia. Molecular Ecology 18, 3683–3698.

58. Guzik M. T., Cooper S. J. B., Humphreys W. F., Ong S., Kawakami T., Austin A. D. (2011b) Evidence for population fragmentation within a subterranean aquatic habitat in the Western Australian desert. Heredity 107(3), 215.

59. Guzik, M. T., Stringer, D. N., Thornhill, J., Coates, P.J., van der Heyde, M., Hillyer, M. J., White, N. E., Saccò, M., Beasley-Hall, P., Humphreys, W. F., Harvey, M. S., Huey, J. A., Wilson, N. G., Alexander, J., Humphreys, G., King, R. A., Cooper, S. J. B., Pinder, A., Perina, G., Nevill, P. and Austin, A. D. (2024) What are the best practices for curating eDNA custom barcode reference libraries? A case study using Australian subterranean fauna. https://biorxiv.org/cgi/content/short/2024.09.18.611555v1.

60. Hajibabaei M., Singer G. A., Hebert P. D., Hickey D. A. (2007) DNA barcoding: how it complements taxonomy, molecular phylogenetics and population genetics. Trends in Genetics 23(4), 167–72.

61. Halse, S.A., (2018) Subterranean fauna of the arid zone. On the Ecology of Australia’s Arid Zone, pp.215–241.

62. Halse S. A., Scanlon M. D., Cocking J. S., Barron H. J., Richardson J. B., Eberhard S. M. (2014) Pilbara stygofauna: deep groundwater of an arid landscape contains globally significant radiation of biodiversity. Records of the Western Australian Museum(2), 443–483.

63. Harvey, F. S., Framenau, V. W., Wojcieszek, J. M., Rix, M. G., & Harvey, M. S. (2012) Molecular and morphological characterisation of new species in the trapdoor spider genus Aname (Araneae: Mygalomorphae: Nemesiidae) from the Pilbara bioregion of Western Australia. Zootaxa, 3383(1), 15–38.

64. Harvey M. S. (1992) The Schizomida (Chelicerata) of Australia. Invertebrate Systematics 6(1), 77–129.

65. Harvey M. S., Berry O., Edward K. L., Humphreys G. (2008) Molecular and morphological systematics of hypogean schizomids (Schizomida:Hubbardiidae) in semiarid Australia. Invertebrate Systematics 22, 167–194.

66. Hassenrück C., Poprick T., Helfer V., Molari M., Meyer R., Kostadinov I. (2021) FAIR enough? A perspective on the status of nucleotide sequence data and metadata on public archives. BioRxiv, 2021.09. 23.461561.

67. Hebert P. D., Cywinska A., Ball S. L., deWaard J. R. (2003) Biological identifications through DNA barcodes. *Proceedings of the Royal Society of London*, Series B 270, 313–322.

68. Hebert P. D., Dewaard J. R., Zakharov E. V., Prosser S. W., Sones J. E., McKeown J. T., Mantle B., La Salle J. (2013) A DNA ‘barcode blitz’: rapid digitization and sequencing of a natural history collection. PLOS ONE 8(7), e68535.

69. Hebert P. D., Gregory T. R. (2005) The promise of DNA barcoding for taxonomy. Systematic Biology 54(5), 852–9.

70. Hlebec D., Podnar M., Kučinić M., Harms D. (2023) Molecular analyses of pseudoscorpions in a subterranean biodiversity hotspot reveal cryptic diversity and microendemism. Scientific Reports 13(1), 430.

71. Jarman S. N., Elliot N. G. (2000) DNA evidence for morphological and cryptic Cenozoic speciations in the Anaspididae, ‘living fossils’ from the Triassic. Journal of Evolutionary Biology 13, 624–633.

72. Javidkar M., Cooper S. J., King R. A., Humphreys W. F., Austin A. D. (2015) Molecular phylogenetic analyses reveal a new southern hemisphere oniscidean family (Crustacea: Isopoda) with a unique water transport system. Invertebrate Systematics 29(6), 554–577.

73. Johnson S. L., Wright A. H. (2001) Central Pilbara groundwater study. Water and Rivers Commission.

74. Karanovic T., Cooper S. (2011) Third genus of parastenocaridid copepods from Australia supported by molecular evidence (Copepoda, Harpacticoida). In ’Studies on freshwater Copepoda: a volume in honour of Bernard Dussart.’ pp. 293–337. (Brill)

75. Karanovic T., Cooper S. J. (2012) ExPLOSive radiation of the genus Schizopera on a small subterranean island in Western Australia (Copepoda: Harpacticoida): unravelling the cases of cryptic speciation, size differentiation and multiple invasions. Invertebrate Systematics 26(2), 115–192.

76. Karanovic T., Eberhard S., Cooper S. J., Guzik M. T. (2015) Morphological and molecular study of the genus Nitokra (Crustacea, Copepoda, Harpacticoida) in a small palaeochannel in Western Australia. Organisms Diversity & Evolution 15, 65–99.

77. Keck F., Altermatt F. (2023) Management of DNA reference libraries for barcoding and metabarcoding studies with the R package refdb. Molecular Ecology Resources 23(2), 511–518.

78. Kimura M. (1980) A simple method for estimating evolutionary rate of base substitutions through comparative studies of nucleotide sequences. Journal of Molecular Evolution 16, 111–120.

79. King R. A., Bradford T., Austin A. D., Humphreys W. F., Cooper S. J. (2012) Divergent molecular lineages and not-so-cryptic species: the first descriptions of stygobitic chiltoniid amphipods (Talitroidea: Chiltoniidae) from Western Australia. Journal of Crustacean Biology 32(3), 465–488.

80. King R. A., Fagan-Jeffries E. P., Bradford T. M., Stringer D. N., Finston T. L., Halse S. A., Eberhard S. M., Humphreys G., Humphreys B. F., Austin A. D. (2022) Cryptic diversity down under: defining species in the subterranean amphipod genus Nedsia Barnard & Williams, 1995 (Hadzioidea: Eriopisidae) from the Pilbara, Western Australia. Invertebrate Systematics 36(2), 113–159.

81. Koch F., Blum P., Korbel K., Menberg K. (2024) Global overview on groundwater fauna. Ecohydrology 17(1), e2607.

82. Korbel K., Chariton A., Stephenson S., Greenfield P., Hose G. C. (2017) Wells provide a distorted view of life in the aquifer: implications for sampling, monitoring and assessment of groundwater ecosystems. Scientific Reports 7, 40702.

83. Kress W. J., Garcia-Robledo C., Uriarte M., Erickson D. L. (2015) DNA barcodes for ecology, evolution, and conservation. Trends in Ecology and Evolution 30(1), 25–35.

84. Kuzmina M. L., Braukmann T. W. A., Fazekas A. J., Graham S. W., Dewaard S. L., Rodrigues A., Bennett B. A., Dickinson T. A., Saarela J. M., Catling P. M., Newmaster S. G., Percy D. M., Fenneman E., Lauron-Moreau A., Ford B., Gillespie L., Subramanyam R., Whitton J., Jennings L., Metsger D., Warne C. P., Brown A., Sears E., Dewaard J. R., Zakharov E. V., Hebert P. D. N. (2017) Using herbarium-derived DNAs to assemble a large-scale DNA barcode library for the vascular plants of Canada. Applied Plant Science 5(12).

85. Kvist S. (2013) Barcoding in the dark? A critical view of the sufficiency of zoological DNA barcoding databases and a plea for broader integration of taxonomic knowledge. Molecular Phylogenetics and Evolution 69(1), 39–45.

86. Kwong S., Srivathsan A., Meier R. (2012) An update on DNA barcoding: low species coverage and numerous unidentified sequences. Cladistics 28(6), 639–644.

87. Lahaye R., Van der Bank M., Bogarin D., Warner J., Pupulin F., Gigot G., Maurin O., Duthoit S., Barraclough T. G., Savolainen V. (2008) DNA barcoding the floras of biodiversity hotspots. Proceedings of the National Academy of Sciences 105(8), 2923–2928.

88. Larson H. K., Foster R., Humphreys W. F., Stevens M. I. (2013) A new species of the blind cave gudgeon Milyeringa (Pisces: Gobioidei, Eleotridae) from Barrow Island, Western Australia, with a redescription of *M. veritas* Whitley. Zootaxa 3616(2), 135–150.

89. Lejzerowicz F., Esling P., Pillet L., Wilding T. A., Black K. D., Pawlowski J. (2015) High-throughput sequencing and morphology perform equally well for benthic monitoring of marine ecosystems. Scientific Reports 5, 13932.

90. Leys R., Watts C. H. S. (2008) Systematics and evolution of the Australian subterranean hydroporine diving beetles (Dytiscidae), with notes on *Carabhydrus*. Invertebrate Systematics 22, 217–225.

91. Leys R., Watts C. H. S., Cooper S. J. B., Humphreys W. F. (2003) Evolution of subterranean diving beetles (Coleoptera: Dytiscidae: Hydroporini, Bidessini) in the arid zone of Australia. Evolution 57(12), 2819–2834.

92. Mächler E., Salyani A., Walser J.-C., Larsen A., Schaefli B., Altermatt F., Ceperley N. (2021a) Environmental DNA simultaneously informs hydrological and biodiversity characterization of an Alpine catchment. Hydrology and Earth System Sciences 25(2), 735–753.

93. Mächler E., Walser J. C., Altermatt F. (2021b) DecisionCmaking and best practices for taxonomyCfree environmental DNA metabarcoding in biomonitoring using Hill numbers. Molecular Ecology 30(13), 3326–3339.

94. Mammola S., Altermatt F., Alther R., Amorim I. R., Băncilă R. I., Borges P. A., Brad T., Brankovits D., Cardoso P., Cerasoli F. (2024) Perspectives and pitfalls in preserving subterranean biodiversity through protected areas. npj Biodiversity 3(1), 2.

95. Mammola S., Cardoso P., Culver D. C., Deharveng L., Ferreira R. L., Fišer C., Galassi D. M., Griebler C., Halse S., Humphreys W. F. (2019) Scientists’ warning on the conservation of subterranean ecosystems. BioScience 69(8), 641–650.

96. Mammola S., Lunghi E., Bilandžija H., Cardoso P., Grimm V., Schmidt S. I., Hesselberg T., Martínez A. (2021) Collecting ecoCevolutionary data in the dark: Impediments to subterranean research and how to overcome them. Ecology and Evolution 11(11), 5911–5926.

97. Matthews E. F., Abrams K. M., Cooper S. J., Huey J. A., Hillyer M. J., Humphreys W. F., Austin A. D., Guzik M. T. (2020) Scratching the surface of subterranean biodiversity: molecular analysis reveals a diverse and previously unknown fauna of Parabathynellidae (Crustacea: Bathynellacea) from the Pilbara, Western Australia. Molecular Phylogenetics and Evolution 142, 106643.

98. Meier R., Blaimer B. B., Buenaventura E., Hartop E., von Rintelen T., Srivathsan A., Yeo D. (2022) A reCanalysis of the data in Sharkey et al.’s (2021) minimalist revision reveals that BINs do not deserve names, but BOLD Systems needs a stronger commitment to open science. Cladistics 38(2), 264–275.

99. Meierotto S., Sharkey M. J., Janzen D. H., Hallwachs W., Hebert P. D., Chapman E. G., Smith M. A. (2019) A revolutionary protocol to describe understudied hyperdiverse taxa and overcome the taxonomic impediment. Deutsche Entomologische Zeitschrift 66(2), 119–145.

100. Meiklejohn K. A., Damaso N., Robertson J. M. (2019) Assessment of BOLD and GenBank– Their accuracy and reliability for the identification of biological materials. PLOS ONE 14(6), e0217084.

101. Mokany K., Harwood T. D., Ferrier S. (2019a) Improving links between environmental accounting and scenarioCbased cumulative impact assessment for betterCinformed biodiversity decisions. Journal of Applied Ecology 56(12), 2732–2741.

102. Mokany K., Harwood T. D., Halse S. A., Ferrier S. (2019b) Riddles in the dark: Assessing diversity patterns for cryptic subterranean fauna of the Pilbara. Diversity and Distributions 25(2), 240–254.

103. Moore G. I., Humphreys W. F., Foster R. (2018) New populations of the rare subterranean blind cave eel Ophisternon candidum (Synbranchidae) reveal recent historical connections throughout north-western Australia. Marine and Freshwater Research 69(10).

104. Mousavi-Derazmahalleh M., Stott A., Lines R., Peverley G., Nester G., Simpson T., Zawierta M., De La Pierre M., Bunce M., Christophersen C. T. (2021) eDNAFlow, an automated, reproducible and scalable workflow for analysis of environmental DNA sequences exploiting Nextflow and Singularity. Molecular Ecology Resources 21(5), 1697–1704.

105. Murphy N. P., Adams M., Guzik M. T., Austin A. D. (2013) Extraordinary micro-endemism in Australian desert spring amphipods. Molecular Phylogenetics and Evolution 66 (3), 645–653.

106. Murphy N. P., Breed M. F., Guzik M. T., Cooper S. J. B., Austin A. D. (2012) Trapped in desert springs: phylogeography of Australian desert spring snails. Journal of Biogeography 39 (9), 1573–1582.

107. Murphy N. P., King R. A., Delean S. (2015) Species, ESUs or populations? Delimiting and describing morphologically cryptic diversity in Australian desert spring amphipods. Invertebrate Systematics 29(5), 457–467.

108. Niemiller M. L., Porter M. L., Keany J., Gilbert H., Fong D. W., Culver D. C., Hobson C. S., Kendall K. D., Davis M. A., Taylor S. J. (2018) Evaluation of eDNA for groundwater invertebrate detection and monitoring: a case study with endangered Stygobromus (Amphipoda: Crangonyctidae). Conservation Genetics Resources 10(2), 247–257.

109. Niemiller, M.L., Helf, K. and Toomey, R.S. (2021) Mammoth cave: a hotspot of subterranean biodiversity in the United States. Diversity, 13(8), 373.

110. Page T. J., Humphreys W. F., Hughes J. M. (2008) Shrimps down under: Evolutionary relationships of subterranean crustaceans from Western Australia (Decapoda: Atyidae: Stygiocaris). PLOS ONE 3(2), 1932–6203.

111. Page T. J., von Rintelen K., Hughes J. M. (2007) Phylogenetic and biogeographic relationships of subterranean and surface genera of Australian Atyidae (Crustacea:Decapoda:Caridea) inferred with mitochondrial DNA. Invertebrate Systematics 21, 137–145.

112. Perina G., Camacho A., Cooper S. J., Floeckner S., Blyth A. J., Saccò M. (2023) An integrated approach to explore the monophyletic status of the cosmopolitan genus Hexabathynella (Crustacea, Bathynellacea, Parabathynellidae): two new species from Rottnest Island (Wadjemup), Western Australia. Systematics and Biodiversity 21(1), 2151662.

113. Perina G., Camacho A., Huey J., Horwitz P., Koenders A. (2018) Understanding subterranean variability: the first genus of Bathynellidae (Bathynellacea, Crustacea) from Western Australia described through a morphological and multigene approach. Invertebrate Systematics 32(2), 423–447.

114. Perina G., Camacho A. I., Huey J., Horwitz P., Koenders A. (2019) New Bathynellidae (Crustacea) taxa and their relationships in the Fortescue catchment aquifers of the Pilbara region, Western Australia. Systematics and Biodiversity 17(2), 148–164.

115. Pochon X., Bott N. J., Smith K. F., Wood S. A. (2013) Evaluating detection limits of next-generation sequencing for the surveillance and monitoring of international marine pests. PLOS ONE 8(9), e73935.

116. Poore G. C. B., Humphreys W. F. (1998) First record of Spelaeogriphacea from Australasia: a new genus and species from an aquifer in the arid Pilbara of Western Australia. Crustaceana 71, 721–742.

117. QGIS.org (2024). QGIS Geographic Information System. Open Source Geospatial Foundation Project. http://qgis.org.

118. Ratnasingham S. (2019) mBRAVE: The Multiplex Barcode Research And Visualization Environment. Biodiversity Information Science and Standards(2).

119. Ratnasingham S., Hebert P. D. (2007) BOLD: The Barcode of Life Data System (http://www.barcodinglife.org). Molecular Ecology Notes 7(3), 355–364.

120. Ratnasingham S., Hebert P. D. N. (2013) A DNA-based registry for all animal species: the Barcode Index Number (BIN) system. PLOS ONE 8(e66213).

121. Rees H. C., Gough K. C., Middleditch D. J., Patmore J. R. M., Maddison B. C., Crispo E. (2015) Applications and limitations of measuring environmental DNA as indicators of the presence of aquatic animals. Journal of Applied Ecology 52(4), 827–831.

122. Rendoš M., Parimuchová A., Hřívová D. K., Karpowicz M., Papáč V., Jabłońska A., Płóciennik M., Haviarová D., Grabowski M. (2023) First insight into molecular diversity and DNA barcode library of epikarst-dwelling invertebrates in the Western Carpathians. Ecohydrology & Hydrobiology.

123. Rideout J. R., He Y., Navas-Molina J. A., Walters W. A., Ursell L. K., Gibbons S. M., Chase J., McDonald D., Gonzalez A., Robbins-Pianka A. (2014) Subsampled open-reference clustering creates consistent, comprehensive OTU definitions and scales to billions of sequences. PeerJ 2, e545.

124. Ruppert K. M., Kline R. J., Rahman M. S. (2019) Past, present, and future perspectives of environmental DNA (eDNA) metabarcoding: A systematic review in methods, monitoring, and applications of global eDNA. Global Ecology and Conservation 17, e00547.

125. Saccò M., Blyth A. J., Douglas G., Humphreys W. F., Hose G. C., Davis J., Guzik M. T., Martínez A., Eberhard S. M., Halse S. A. (2022a) Stygofaunal diversity and ecological sustainability of coastal groundwater ecosystems in a changing climate: The Australian paradigm. Freshwater Biology 67(12), 2007–2023.

126. Saccò M., Guzik M. T., van der Heyde M., Nevill P., Cooper S. J., Austin A. D., Coates P. J., Allentoft M. E., White N. E. (2022b) eDNA in subterranean ecosystems: Applications, technical aspects, and future prospects. Science of The Total Environment, 153223.

127. Saccò M., Mammola S., Altermatt F., Alther R., Bolpagni R., Brancelj A., Brankovits D., Fišer C., Gerovasileiou V., Griebler C. (2024) Groundwater is a hidden global keystone ecosystem. Global Change Biology 30(1), e17066.

128. Saclier N., Duchemin L., Konecny-Dupré L., Grison P., Eme D., Martin C., Callou C., Lefébure T., François C., Issartel C., Lewis J. J., Stoch F., Sket B., Gottstein S., Delić T., Zagmajster M., Grabowski M., Weber D., Reboleira A. S. P. S., Palatov D., Paragamian K., Knight L. R. F. D., Michel G., Lefebvre F., Hosseini M.-J. M., Camacho A. I., De Bikuña B. G., Taleb A., Belaidi N., Tuekam Kayo R. P., Galassi D. M. P., Moldovan O. T., Douady C. J., Malard F. (2024) A collaborative backbone resource for comparative studies of subterranean evolution: The World Asellidae database. Molecular Ecology Resources 24(1), e13882.

129. Saitou N., Nei M. (1987) The neighbor-joining method: a new method for reconstructing phylogenetic trees. Molecular Biology and Evolution 4(4), 406–425.

130. Sala (2000) Global biodiversity Scenarios for the year 2100. Science.

131. Simon C., Frati F., Beckenbach A., Crespi B., Liu H., Flook P. (1994) Evolution, weighting, and phylogenetic utility of mitochondrial gene sequences and a compilation of conserved polymerase chain reaction primers. Annals of the Entomological Society of America 87(6), 651–701.

132. Sket, B., Paragamian, K., Tontelj, P. (2004) A census of the obligate subterranean fauna of the Balkan Peninsula. In: Griffith HI (ed) Balkan biodiversity. Kluwer Academic, Dordrecht, pp 309–322

133. Stall S., Yarmey L., Cutcher-Gershenfeld J., Hanson B., Lehnert K., Nosek B., Parsons M., Robinson E., Wyborn L. (2019) Make scientific data FAIR. Nature 570(7759), 27–29.

134. Stringer D. N., King R. A., Austin A. D., Guzik M. T. (2022) Pilbarana, a new subterranean amphipod genus (Hadzioidea: Eriopisidae) of environmental assessment importance from the Pilbara, Western Australia.

135. Takahara, T., Minamoto, T., Yamanaka, H., Doi, H., & Kawabata, Z. I. (2012). Estimation of fish biomass using environmental DNA. PLOS one 7(4), e35868.

136. Takahashi M., Saccò M., Kestel J. H., Nester G., Campbell M. A., Van Der Heyde M., Heydenrych M. J., Juszkiewicz D. J., Nevill P., Dawkins K. L. (2023) Aquatic environmental DNA: A review of the macro-organismal biomonitoring revolution. Science of The Total Environment 873, 162322.

137. Thomsen P. F., Kielgast J., Iversen L. L., Wiuf C., Rasmussen M., Gilbert M. T., Orlando L., Willerslev E. (2012) Monitoring endangered freshwater biodiversity using environmental DNA. Molecular Ecology 21(11), 2565–73.

138. Thomsen P. F., Willerslev E. (2015) Environmental DNA – An emerging tool in conservation for monitoring past and present biodiversity. Biological Conservation 183, 4–18.

139. Tixier M.-S., Hernandes F. A., Guichou S., Kreiter S. (2012) The puzzle of DNA sequences of Phytoseiidae (Acari: Mesostigmata) in the public GenBank database. Invertebrate Systematics 25(5), 389–406.

140. Vamos E., Elbrecht V., Leese F. (2017) Short COI markers for freshwater macroinvertebrate metabarcoding. Metabarcoding and Metagenomics 1.

141. van der Heyde M., Alexander J., Nevill P., Austin A. D., Stevens N., Jones M., Guzik M. T. (2023a) Rapid detection of subterranean fauna from passive sampling of groundwater eDNA. Environmental DNA 5(6), 1706–1719.

142. van der Heyde M., White N. E., Nevill P., Austin A. D., Stevens N., Jones M., Guzik M. T. (2023b) Taking eDNA underground: factors affecting eDNA detection of subterranean fauna in groundwater. Molecular Ecology Resources.

143. Vilgalys R. (2003) Taxonomic misidentification in public DNA databases. New Phytologist 160(1), 4–5.

144. Webb J. M., Jacobus L. M., Funk D. H., Zhou X., Kondratieff B., Geraci C. J., DeWalt R. E., Baird D. J., Richard B., Phillips I., Hebert P. D. (2012) A DNA barcode library for North American Ephemeroptera: progress and prospects. PLOS ONE 7(5), e38063.

145. White N. E., Guzik M. T., Austin A. D., Moore G. I., Humphreys W. F., Alexander J., Bunce M. (2020) Detection of the rare Australian endemic blind cave eel (Ophisternon candidum) with environmental DNA: implications for threatened species management in subterranean environments. Hydrobiologia 847, 3201–3211.

146. Wilkinson M. D., Dumontier M., Aalbersberg I. J., Appleton G., Axton M., Baak A., Blomberg N., Boiten J.-W., da Silva Santos L. B., Bourne P. E. (2016) The FAIR Guiding Principles for scientific data management and stewardship. Scientific data 3(1), 1–9.

147. Wilkinson S. P., Davy S. K., Bunce M., Stat M. (2024) Taxonomic identification of environmental DNA with informatic sequence classification trees. PeerJ 12, e16963.

148. Wirta H., Varkonyi G., Rasmussen C., Kaartinen R., Schmidt N. M., Hebert P. D., Bartak M., Blagoev G., Disney H., Ertl S., Gjelstrup P., Gwiazdowicz D. J., Hulden L., Ilmonen J., Jakovlev J., Jaschhof M., Kahanpaa J., Kankaanpaa T., Krogh P. H., Labbee R., Lettner C., Michelsen V., Nielsen S. A., Nielsen T. R., Paasivirta L., Pedersen S., Pohjoismaki J., Salmela J., Vilkamaa P., Vare H., von Tschirnhaus M., Roslin T. (2016) Establishing a community-wide DNA barcode library as a new tool for arctic research. Molecular Ecology Resources 16(3), 809–22.

149. Zahiri R., Lafontaine J. D., Schmidt B. C., Dewaard J. R., Zakharov E. V., Hebert P. D. (2017) Probing planetary biodiversity with DNA barcodes: The Noctuoidea of North America. PLOS ONE 12(6), e0178548.

150. Zhou X., Adamowicz S. J., Jacobus L. M., Dewalt R. E., Hebert P. D. (2009) Towards a comprehensive barcode library for arctic life - Ephemeroptera, Plecoptera, and Trichoptera of Churchill, Manitoba, Canada. Frontiers in Zoology 6, 30.

