## Supplementary Material 1 for "Towards a global barcode reference library for subterranean fauna"

**Supplementary Material 1:** Global map showing single exemplar subterranean biodiversity hotspot locations per continent, their respective major taxon group diversity of troglofauna and stygofauna, the number of species and corresponding references for original data.

| **Continent** | **Region/Location** | **Stygofauna taxon groups** | **Troglofauna taxon groups** | **Stygofauna** | **Troglofauna** | **Reference** |
| --- | --- | --- | --- | --- | --- | --- |
| Asia | Hon Chong karst, Vietnam | **Crustacea:**  Amphipoda  Isopoda  **Mollusca:**  Gastropoda | **Arachnida**:  Acarina  Araneae  Opiliones  Pseudoscorpiones  **Crustacea:**  Isopoda  **Hexapoda**:  Blattodea  Collembola  Coleoptera  Diplura  Hemiptera  Zygentoma  **Myriapoda:**  Diplopoda | 3 | 36 | Deharveng *et al.* (2023) |
| Australia | Pilbara, Western Australia | **Arachnida:**  Acarina  **Annelida**  **Crustacea:**  Amphipoda  Copepoda  Isopoda  Ostracoda  Syncarida  Thermosbaenacea  **Mollusca:**  Gastropoda  **Nematoda**  **Platyhelminthes**  **Rotifera**  **Vertebrata:**  Actinopterygii (eel) | **Arachnida**:  Pseudoscorpiones  Schizomida  Araneae  Palpigradi  **Crustacea:**  Isopoda  **Hexapoda:**  Blattodea  Coleoptera  Diplura  Hemiptera  Zygentoma  **Myriapoda**:  Chilopoda  Diplopoda  Pauropoda  Symphyla | 637 | 680 | Halse (2018) |
| Europe | Balkan Peninsula | **Annelida**  **Cnidaria (jellyfish)**  **Crustacea:**  Amphipoda  Cladocera (water flea)  Copepoda  Decapoda  Isopoda  Mysidacea  Ostracoda  Syncarida  Thermosbaenacea  **Mollusca:**  Gastropoda  Bivalvia  **Nemertea (ribbon worm)**  **Platyhelminthes**  **Porifera (sponge)**  **Vertebrata**:  Amphibia | **Arachnida**:  Araneae  Opiliones  Palpigradi  Pseudoscorpiones  **Crustacea**:  Isopoda  **Hexapoda**:  Collembola  Coleoptera  Diplura  Diptera  Zygentoma  **Mollusca:**  Gastropoda  **Myriapoda**:  Chilopoda,  Diplopoda  **Platyhelminthes** | 686 | 995 | Sket *et al.* (2004) |
| Africa | Wynberg cave system, South Africa | **Crustacea:**  Amphipoda  Spelaeogriphacea | **Arachnida**:  Acarina  Araneae  Opiliones  Pseudoscorpiones  **Crustacea:**  Isopoda  **Hexapoda**:  Collembola  Coleoptera  Diplura  **Myriapoda:**  Diplopoda  **Onychophora**  **Platyhelminthes** | 2 | 17 | Ferreira *et al.* (2020) |
| North America | Mammoth Cave, Kentucky, U.S. A. | **Crustacea:**  Amphipoda  Copepoda  Decapoda  Isopoda  Ostracoda  **Mollusca:**  Gastropoda  **Vertebrata:**  Actinopterygii (fish) | **Arachnida:**  Acarina  Araneae  Opiliones  Pseudoscorpiones  **Hexapoda:**  Collembola  Coleoptera  Diplura  Diptera  Psocodea  **Mollusca:**  Gastropoda  **Myriapoda:**  Diplopoda  **Platyhelminthes** | 17 | 32 | Niemiller *et al.* (2021) |
| South America | Chapada Diamantina Region, Brazil | **Vertebrata:**  Actinopterygii (fish) | **Arachnida:**  Acarina  Araneae  Opiliones  Palpigradi  Pseudoscorpiones  Scorpiones  **Crustacea:**  Isopoda  **Hexapoda:**  Blattodea  Collembola  Coleoptera  Diplura  Zygentoma  **Mollusca:**  Gastropoda  **Myriapoda**:  Chilopoda,  Diplopoda | 2 | 35 | Gallão *et al.* (2023) |

**REFERENCES**

Deharveng, L., Le, C.K., Bedos, A., Judson, M.L., Le, C.M., Lukić, M., Luu, H.T., Ly, N.S., Nguyen, T.Q.T., Truong, Q.T. and Vermeulen, J. (2023) A hotspot of subterranean biodiversity on the brink: Mo So Cave and the Hon Chong Karst of Vietnam. *Diversity* **15**(10), 1058.

Ferreira, R.L., Giribet, G., Du Preez, G., Ventouras, O., Janion, C. and Silva, M.S. (2020) The Wynberg cave system, the most important site for cave fauna in South Africa at risk. *Subterranean Biology* **36**, 73-81.

Gallão, J.E., Ribeiro, D.B., Gallo, J.S. and Bichuette, M.E., 2023. There and Back Again—The Igatu Hotspot Siliciclastic Caves: Expanding the data for subterranean fauna in Brazil, Chapada Diamantina Region. *Diversity* **15**(9), 991.

Halse, S.A., (2018) Subterranean fauna of the arid zone. On the Ecology of Australia’s Arid Zone, pp.215-241.

Niemiller, M.L., Helf, K. and Toomey, R.S. (2021) Mammoth cave: a hotspot of subterranean biodiversity in the United States. *Diversity*, **13**(8), 373.

Sket, B., Paragamian, K., Tontelj, P. (2004) A census of the obligate subterranean fauna of the Balkan Peninsula. In: Griffith HI (ed) Balkan biodiversity. Kluwer Academic, Dordrecht, pp 309–322.
